# Complexes of tubulin oligomers and tau form an intervening network cross-bridging microtubules into bundles

**DOI:** 10.1101/2022.07.18.500499

**Authors:** Phillip Kohl, Chaeyeon Song, Bretton Fletcher, Rebecca L. Best, Christine Tchounwou, Ximena Garcia Arceo, Peter J. Chung, Herbert P. Miller, Leslie Wilson, Myung Chul Choi, Youli Li, Stuart C. Feinstein, Cyrus R. Safinya

**Affiliations:** Materials Research Laboratory, University of California, Santa Barbara; Santa Barbara, CA 93106, USA; Materials Department, University of California, Santa Barbara; Santa Barbara, CA 93106, USA; Biomolecular Science and Engineering, University of California, Santa Barbara; Santa Barbara, CA 93106, USA; Department of Molecular, Cellular, and Developmental Biology, University of California, Santa Barbara; Santa Barbara, CA, USA; Department of Physics, University of California, Santa Barbara; Santa Barbara, CA 93106, USA; Neuroscience Research Institute, University of California, Santa Barbara; Santa Barbara, CA 93106, USA; Department of Bio and Brain Engineering, Korea Advanced Institute of Science and Technology, 291 Daehak-ro, Daejeon 34141, Korea

**Keywords:** tubulin oligomers, tau, intrinsically disordered proteins, microtubule bundles, microtubule fascicles, phase diagram, SAXS, plastic embedded TEM

## Abstract

The axon-initial-segment (AIS) of mature neurons contains microtubule (MT) fascicles (linear bundles) that are implicated as retrograde diffusion barriers in the retention of MT-associated protein (MAP) tau inside axons. While the role of tau in MT bundling is poorly understood, tau dysfunction and leakage outside of the axon is associated with neurodegeneration. We report on the structure of steady-state MT bundles in response to varying concentrations of divalent cations (Mg^2+^ or Ca^2+^) in dissipative reaction mixtures containing αβ-tubulin, full-length tau, and GTP at 37°C. A concentration-time kinetic phase diagram generated by synchrotron small-angle X-ray scattering (SAXS) reveals a wide-spacing MT bundle phase (B_ws_), a transient intermediate MT bundle phase (B_int_), and a tubulin ring phase. Remarkably, SAXS analysis combined with TEM of plastic embedded samples provides direct evidence of an intervening network (IN) of complexes of tubulin oligomers and tau (≈5 nm wide filaments), which stabilize MT bundles. In this model, αβ-tubulin oligomers in the IN are crosslinked by tau’s MT binding repeats, which also link αβ-tubulin oligomers to αβ-tubulin within the MT lattice. The finding of a new role for tubulin revises current dogma where cross-bridging of MTs is attributed entirely to interactions between MAPs. The tubulin-tau complexes of the IN should enhance the barrier properties of MT fascicles in preventing tau missorting to the somatodendritic compartment as happens during neurodegeneration. Furthermore, tubulin-tau complexes in the IN or bound to isolated MTs are potential sites for enzymatic modification of tau promoting nucleation and growth of tau fibrils in tauopathies.

**Significance Statement:** A cell free model of microtubule (MT) bundles of the axon-initial-segment (known as MT fascicles) was studied in physiologically relevant buffer conditions. MT fascicles have a role in retaining neuronal protein tau, a key protein stabilizing MTs, in the axon. X-ray scattering and electron microscopy led to the discovery of complexes of tubulin oligomers and tau as building blocks of an intervening network that cross-bridge MTs into stable bundles with precisely the same linear geometry observed *in-vivo* in neurons. Significantly, changes to the chemical structure of tau because of abnormal interactions with cellular enzymes, would be predicted to disrupt the intervening tubulin-tau network and the MT-fascicle’s barrier function, promoting leakage of tau to the somatodendritic compartment and neuron degradation.

---

Microtubules (MTs) are hollow protein nanotubes resulting from GTP-mediated assembly of αβ-tubulin heterodimers, which stack to form protofilaments (PFs, tubulin oligomers) with lateral PF-PF interactions stabilizing the MT wall (*1*). Upon hydrolysis of GTP in the β-tubulin subunit, GDP-PFs transition to a higher curvature conformation. These distinct PF conformations underlie MT dynamic instability (DI), corresponding to stochastic switching between periods of slow growth (polymerization) and rapid depolymerization (2-6). DI is regulated by the ratio of GTP- to GDP-tubulin where high concentrations of GTP-tubulin stabilize a GTP-tubulin cap promoting MT polymerization, while low concentrations lead to loss of the cap and subsequent inside-out curving of GDP-PFs and rapid MT disassembly (*7*). In cells, MT-associated proteins (MAPs) regulate DI and are implicated in MT bundle formation (*8-12*). Dynamic MT bundles are involved in various cell functions including chromosome segregation (*9*) and axonogenesis in developing neurons (*13, 14*). Linear MT bundles with large wall-to-wall spacing (referred to as fascicles of MTs, SI Appendix, Fig. S1), found in the central core of the axon initial segment (AIS) of mature neurons (*15-17*), form a retrograde diffusion barrier of MAP tau (*18*) with a role in the compartmentalization of tau in axons, as well as axonal transport.

In our study, we focused on mixtures of αβ-tubulin and the MAP tau, an intrinsically disordered protein, which upon binding to MTs partially suppresses DI (*19-22*) (*23, 24*) and facilitates the transport of cargo along MTs in axons (*9*). Tau dysfunction is implicated in neurodegenerative “tauopathies”, which include Alzheimer’s disease (*25*), FTDP-17 (*26*), and chronic traumatic encephalopathy (*27*). Humans express six wild-type tau isoforms resulting from alternative splicing of exons 2, 3, and 10 of the MAPT gene (Fig. 1A) (*28*). The N-terminal projection domain (PD) of tau (protruding away from the MT surface) is followed by a proline-rich region, the MT binding region (MTBR), and a carboxyl-terminal tail (*29-31*). We used full length (4RL) tau where the MTBR includes four (R1-R4) 18 amino acid imperfect repeat sequences separated by 13 to 14 amino acid inter-repeats, and the N-terminal tail includes inserts 1 (N1) and 2 (N2) encoded by exons 2 and 3, respectively (Fig. 1A). Tau binding to the negatively charged carboxyl terminal residues of αβ-tubulin is facilitated by the cationic nature of the MTBR and regions flanking the MTBR, in particular, the proline-rich region.

**Figure 1.**
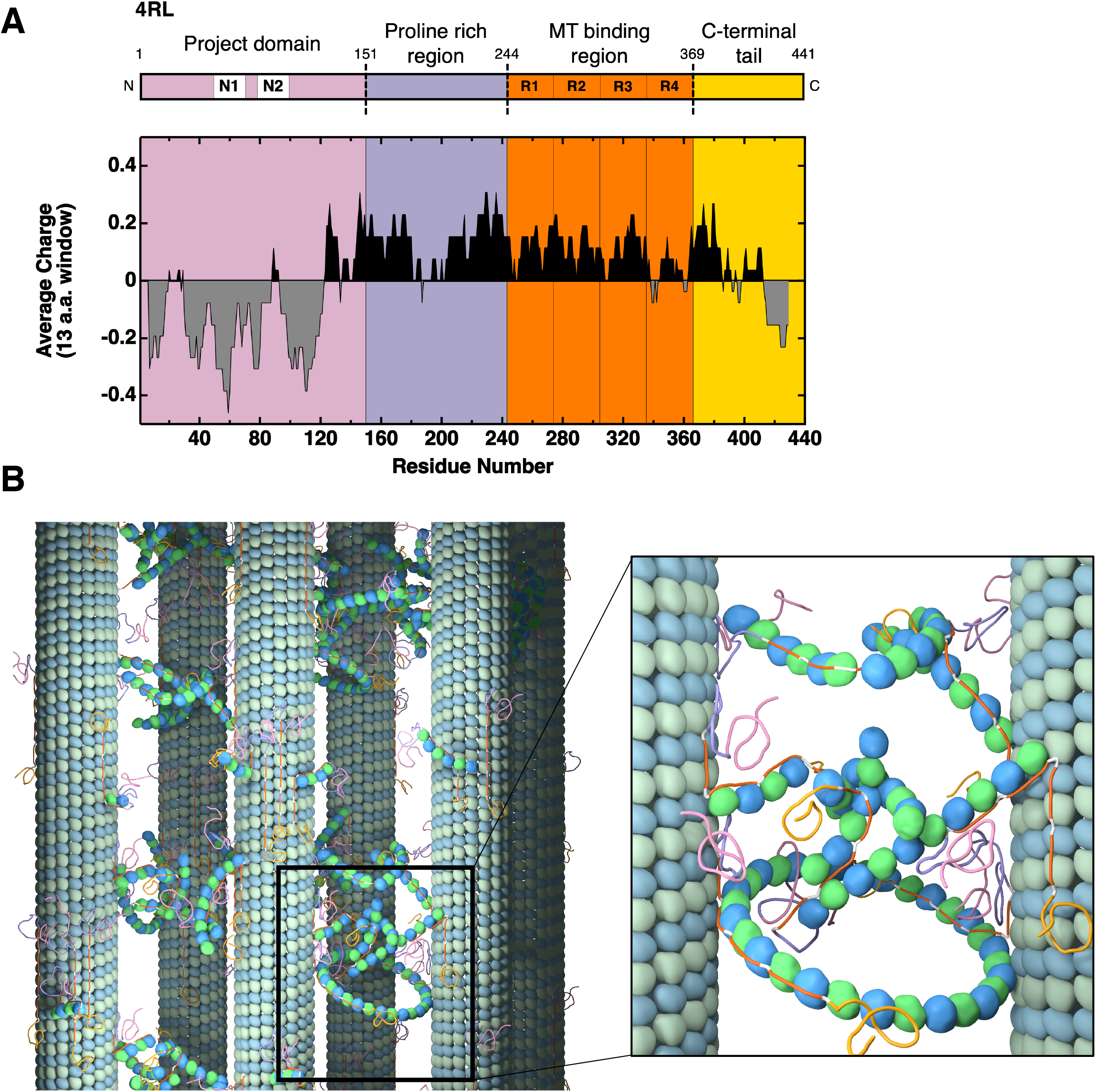
Tau charge distribution and tau-tubulin network stabilizing bundled MTs. (**A**) Schematic and average charge profile of full length 4RL tau with major features labeled, including inserts 1 and 2, encoded by exons 2 and 3, respectively, and the MT-binding repeats (R1-R4). 4RL tau is depicted with labeled domains: N-terminal tail consisting of the projection domain (PD) and proline rich region, the microtubule (MT) binding region (MTBR), and the C-terminal tail (*29-31*). The charge distribution is calculated using a rolling sum over thirteen residues. Despite anionic regions at the N- and C-termini, tau isoforms have a net cationic charge, which contributes to tau’s binding (via MTBR) to negatively charged residues of αβ-tubulin. Data are from the National Center for Biotechnology Information Protein Database (accession number NP_005901.2). (**B**) Cartoon of a microtubule bundle (left) with blow-up (right) showing an intervening network of complexes of tubulin oligomers (curved short protofilaments and rings) and tau, which stabilizes MT bundles in the B_ws_ and B_int_ phases. The cationic MT binding repeats of tau (orange sections) link αβ-tubulin oligomers both to other free tubulin oligomers and to αβ-tubulin dimers within the MT lattice, creating the intervening network that cross-bridges neighboring MTs. Tau is depicted to bind either side of curved tubulin oligomers and rings consistent with studies that show tau may bind (via the MT binding repeats) either the outside surface or lumen of the MT(*37, 64, 65*).

The current study was designed to elucidate the structure and stability of MT bundles in response to increasing concentrations of divalent cations (Mg^2+^ or Ca^2+^) in minimal dissipative reaction mixtures at 37°C containing αβ-tubulin, 4RL-tau, and GTP in standard buffer (*32*). The mixtures contained a 1/20 tau/tubulin-dimer molar ratio, which corresponds to sub-monolayer coverage for tau on MTs in the mushroom regime (*33*). The divalent cation concentrations were in the millimolar range, which for Mg^2+^ approximates average cell concentrations (*34*). Thus, a motivation for the studies with physiological concentrations of Mg^2+^ was that they represented a minimal cell free model of MT fascicles of the AIS.

Time-dependent synchrotron small-angle X-ray scattering (SAXS) was used to monitor samples over 33 hours and revealed three distinct states of tubulin structures, generating a kinetic phase diagram of tau-tubulin structures as a function of time and concentrations of Mg^2+^ or Ca^2+^. The phase diagram guided the selection of specific time points for parallel TEM of plastic embedded samples, for real-space images and comparisons to reciprocal-space SAXS data. In agreement with previous measurements by Chung et al. (*35*), widely-spaced MT bundles (labeled B_ws_ with MT wall-to-wall spacing *d*_w-w_ ≈ 40 to 45 nm) were found to be stable below a critical lower divalent cation concentration c=c_lower_ (≈0.8 mM CaCl_2_ and ≈1.6 mM MgCl_2_).

Over a narrow range of Ca^2+^ or Mg^2+^ concentrations (c_lower_<c<c_upper_ c_upper_ ≈1.4 mM CaCl_2_ and ≈2.4 mM MgCl_2_), the B_ws_ state evolves over time to a tubulin ring state. The structural evolution, which is signaled by the depolymerization of a fraction of MTs, leads to a sudden increase in the formation of tubulin rings and curved tubulin oligomers (observed in SAXS), which occurs over a period of order hours. During this time period, the newly created tubulin oligomers (observed in TEM as ≈5 nm wide flexible filaments and rings) are found in between MTs in bundles leading to the creation of a transient intermediate bundle state (labeled B_int_) with a more ordered lattice with smaller *d*_w-w_ (≈30-35 nm). This is observed in TEM by the presence of significantly more cross-bridges between MTs in the B_int_ compared to the B_ws_ state (albeit with similar morphologies).

Taken together, SAXS and TEM data are consistent with MTs bundled by an intervening network consisting of complexes of tubulin oligomers and tau in both the B_ws_ and B_int_ states. In this model, which represents a significant revision to current dogma (*8, 9, 12, 36*), MT bundles are formed due to “coded assembly” where tau’s MT binding repeats (orange sections in Fig. 1A and 1B) are the glue linking αβ-tubulin oligomers in the intervening network. Furthermore, tau also links αβ-tubulin oligomers, in the intervening network near the MT surface, to αβ-tubulin in the MT lattice (Fig. 1B). Cross-bridges between MTs have been reported previously but were attributed solely to MAPs (*8, 10-13*). In our model, the large MT wall-to-wall spacings observed in the bundled states (much larger than the size of tau’s PD) is set by the average radius of curvature of tau-coated curved tubulin oligomers and rings.

## Results

### Time-dependent SAXS generates a kinetic phase diagram revealing three distinct assembly structures for αβ-tubulin/tau/divalent cation/GTP mixtures at 37°C

To understand how divalent cation content modulates the stability and structural features of tau-mediated MT bundles, time-dependent synchrotron SAXS measurements were performed on tubulin reaction mixtures (30 – 40 µM) containing 2 mM GTP co-assembled at 37 °C with 4RL-tau (tau to tubulin-dimer molar ratio, Φ_4RL_ = 0.05) at varying CaCl_2_ or MgCl_2_ concentrations (0 to 5 mM added to buffer). Analysis of azimuthally averaged SAXS profiles at initial time point (*t*_0_) soon after sample preparation, revealed three distinct phases with increasing divalent cation concentration (Fig. 2A).

**Figure 2.**
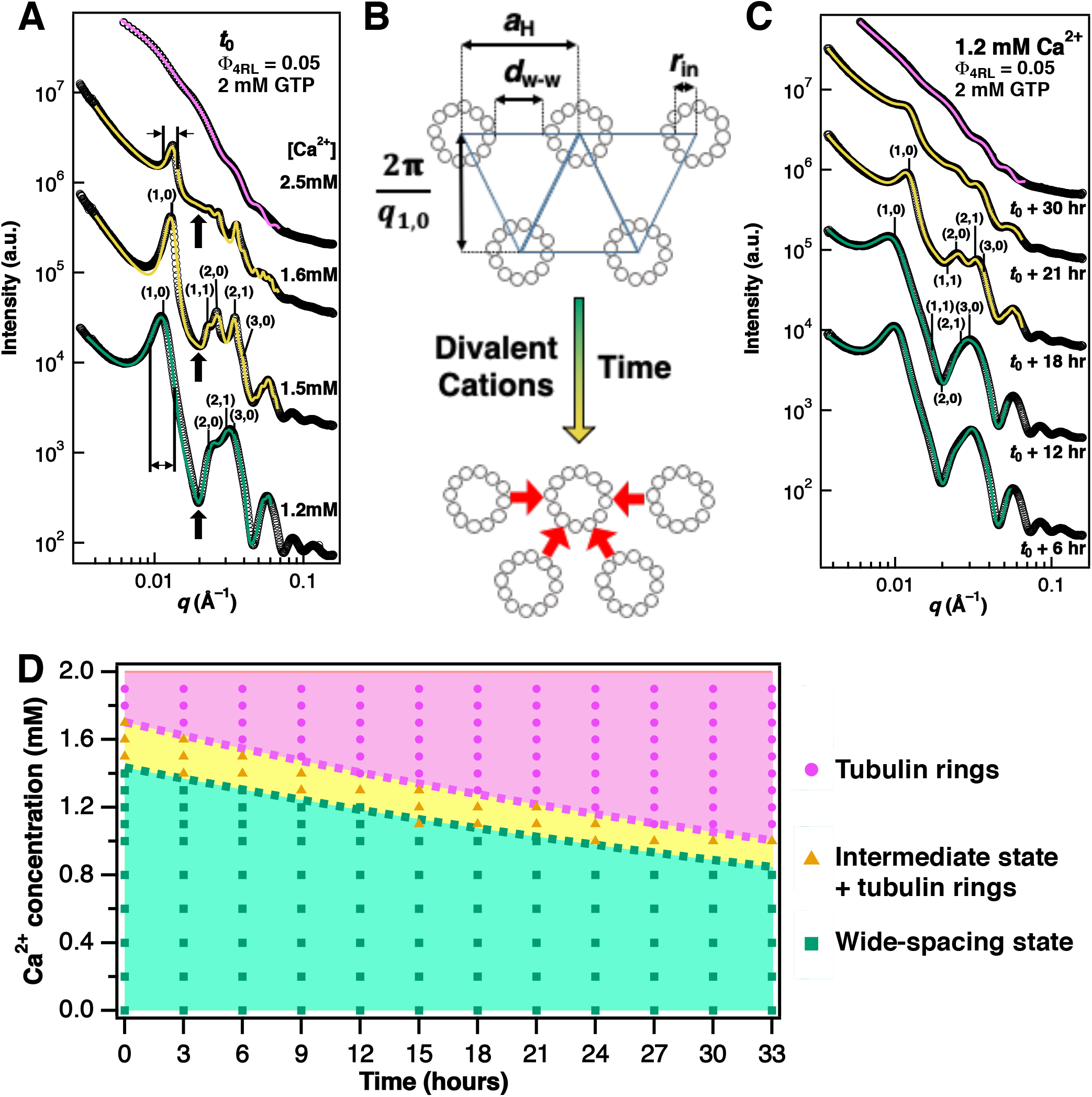
Ca^2+^- and time- dependent synchrotron SAXS data reveals an intermediate bundled (B_int_) microtubule state between the bundled wide-spacing (B_ws_) and the tubulin ring states. (**A**) SAXS data (open circles) and corresponding fits (solid lines) for increased Ca^2+^ concentrations at t_0_, where t_0_ is the time of the first measurement. Indexing for three lower profiles is for 2D hexagonal lattice of MTs. The intermediate bundle state (B_int_, profiles at 1.5mM and 1.6mM Ca^2+^) is differentiated from the widely-spaced bundle state (B_ws_, profile at 1.2 mM Ca^2+^) by peaks shifted to larger q (i.e. smaller lattice spacing), decreased peak widths (compare 1,0 peak positions for 1.6mM and 1.2mM Ca^2+^), and increased scattering intensity at local minima of the MT form factor (solid arrows at 1.2mM, 1.5mM, and 1.6mM Ca^2+^). At 2.5 mM Ca^2+^ the SAXS is dominated by tubulin rings and curved oligomers. (**B**) Cartoon of hexagonal MT bundles in the B_ws_ (top) and B_int_, (bottom) states highlights changes in MT-MT spacing. (**C**) SAXS data (open circles) and corresponding fits (solid lines) with increasing time at 1.2 mM Ca^2+^. Evolution of SAXS profiles show that phase transitions occur from B_ws_ to B_int_ (between t_0_ +12 and t_0_ +18 hrs) and from B_int_ to the tubulin ring state (between t_0_ +18 and t_0_ +30 hrs). (**D**) Kinetic phase diagram of tau/tubulin mixtures as a function of Ca^2+^ concentration and time. Distinct B_ws_, B_int_, and (tau-coated) tubulin ring states are observed. Markers indicate the phase, as determined by SAXS analysis, of each data point and are color-coded to match the line color of the SAXS fits plotted in (A) and (C). While details of the phase diagram (i.e. critical concentrations, the timing of transitions) vary between batches of tubulin and tau, the data represents the repeatable trends for mixtures containing either added CaCl_2_ or MgCl_2_.

All samples below threshold divalent concentrations of 1.4 mM for Ca^2+^ and 2.4 mM for Mg^2+^ exhibited MT bundling characteristics indistinguishable from controls with no added divalent cations (SI Appendix, Fig. S2) and are consistent with reported values of the widely-spaced MT bundle state *(35)*. A typical example of a SAXS profile for a sample in the B_ws_ phase is plotted in Fig. 2A at 1.2 mM Ca^2+^, which displays scattering characteristics indicative of strong MT polymerization and registers Bragg peaks consistent with 2D hexagonal packing of MTs (*q*_10_, *q*_11_=3^1/2^*q*_10_, *q*_20_=2*q*_10_, *q*_21_=7^1/2^*q*_10_, *q*_30_=3*q*_10_, *q*_22_=12^1/2^*q*_10_). The location of the (1,0) peak, *q*_10_, which is used to measure the center-to-center distance between microtubules, *a*_H_ = 4π/(*q*_10_√3) (Fig. 2B), ranged between *q*_10_ = 0.0115 Å^−1^ and 0.0122 Å^−1^ (interaxial MT spacing between *a*_H_ = 59.5 and 63.1 nm).

However, tau-tubulin reaction mixtures above these threshold concentrations produced scattering profiles at *t*_*0*_ that were markedly different and indicative of a new intermediate bundled state (B_int_) (Fig. 2A, profiles at 1.5mM and 1.6mM Ca^2+^). A notable difference between the B_ws_ and B_int_ states (Fig. 2A, 1.2 mM compared to 1.5 mM Ca^2+^) is the characteristic shift in location of all Bragg peaks to higher *q* (*q*_10_ for B_int_ ranged between 0.0128 to 0.133 Å ^−1^, *a*_H_ = 54.5 to 56.7 nm), implying that the interaxial spacing of the B_int_ state is smaller than in the B_ws_ phase (by approximately 7 to10 nm). The Bragg peaks are also sharper in the B_int_ than in the B_ws_ state—as illustrated in Fig. 2A by measurements of the full-width at half maximum (FWHM) for *q*_10_ (Fig. 2A, vertical lines at 1.2 mM and 1.6 mM Ca^2+^). This difference indicates that, despite having a smaller lattice parameter, the coherent domain size of the MT lattice (inversely proportional to the FWHM) is much larger in the B_int_ state. Interestingly, despite the larger domain size of the MT lattice, there is a significant decrease in scattering intensity from bundled MTs for samples in the B_int_ state. This observation is more pronounced with increasing Ca^2+^ concentration (Fig. 2A, scattering intensity of (1,0) peak diminishes with increased Ca^2+^) and indicates that within this phase, Ca^2+^ severely inhibits MT polymerization in a concentration-dependent manner. This effect can be seen unambiguously by noting the increase in scattering at the profile’s local minima with increased cation content (Fig. 2A, arrows) due to the increased prevalence of depolymerized tubulin products (discussed below).

At higher Ca^2+^ concentrations (1.9 - 3.0 mM Ca^2+^), MT polymerization was substantially inhibited and SAXS profiles were indicative of single tubulin rings (Fig. 2A, broad oscillations at 2.5 mM Ca^2+^) and curved tubulin oligomers. SAXS signatures of the tubulin ring state continued to dominate and remained unchanged with increased divalent concentrations. As has been shown previously for free tubulin heterodimers and oligomers (*37-39*) and for tubulin spiral structures under non-assembly promoting conditions(*40*), we expect these curved tubulin structures to be coated with tau via the MT binding repeats.

Intriguingly, the Mg^2+^- or Ca^2+^-induced structural transitions also occur as a function of time for samples with intermediate concentrations of added Ca^2+^ or Mg^2+^ (c_lower_≈0.8 to c_upper_≈1.4 mM Ca^2+^ or c_lower_≈1.2 to c_upper_≈2.2 mM Mg^2+^). These samples originate in the B_ws_ state but abruptly transition to the B_int_ state after several hours. A typical example is shown in Fig. 2C where a sample with 1.2 mM Ca^2+^ is in the B_ws_ state at *t*_0_ +6 hrs and *t*_0_ +12 hrs, the B_int_ phase at *t*_0_ +18 hrs and *t*_0_ +21 hrs, and the ring state at *t*_0_ +30 hrs. The defining characteristics of the B_int_ state compared to the B_ws_ state observed along the concentration axis (smaller lattice spacing, larger bundle domain size) can also be seen here with increasing time. Similarly, the transition to the B_int_ state is accompanied by a significant increase in scattering at the SAXS profile’s first local minima with increased time (Fig. 2C compare profiles at *t*_0_ +6 hrs and *t*_0_ +12 hrs to *t*_0_ +18 hrs and *t*_0_ +21 hrs). This is due to the rapid increase in the rate of MT depolymerization and tubulin ring proliferation, which, as we show later, drives the transition from the B_ws_ to the B_int_ state. Depolymerization of nearly all MTs and the structural evolution to the ring state is typically observed between 6 to 9 hours after the B_ws_ to B_int_ transition.

Figure 2D summarizes the SAXS data in a kinetic phase diagram displaying distinct regions where the B_ws_ (green), B_int_ (yellow), and (tau-coated) tubulin ring (magenta) are the dominant structures as a function of Ca^2+^ concentration and time. The B_int_ state is a two-phase system with bundled MTs and tubulin rings coexisting in amounts readily detectable by SAXS. The start of the region in the phase diagram denoted as the ring state corresponds to the earliest time point at which the fitted q_10_ peak of the B_int_ phase can no longer be resolved. Data reveals a clear decrease in the lifetime of the B_ws_ state with increased divalent ion content, and is likely related to a similar effect by tetra-valent spermine, which promotes depolymerization of paclitaxel (PTX)-stabilized MTs, although on longer time scales of order days at room temperature (*41*) due to the larger stabilizing effect of PTX on MTs compared to tau.

### Plastic-embedded TEM images are consistent with SAXS data and reveal distinct structural features in the B_ws_ and B_int_ bundled phases

To better understand the structural differences between the two bundled states, TEM experiments were performed on plastic-embedded tau/tubulin/2mM GTP reaction mixtures in standard buffer containing 1.8 mM of added Mg^2+^ at 37 °C (SI Appendix, SI Note S2). Samples were individually prepared and fixed 3 hours (Fig. 3 A-C) and 18 hours (Fig. 3 D-F) after polymerization, time points where the wide-spacing B_ws_ and the intermediate B_int_ bundled phases were observed via SAXS in samples containing 1.8 mM added Mg^2+^, respectively (discussed below, Fig. 4C). Consistent with SAXS line-shape analysis, cross-sectional images at lower (Fig. 3 A,D) and higher (Fig. 3 B,E) magnification show clear phase separation into distinct bundled domains at both time points. Larger domain sizes (bundle widths) are seen at 18 hours (Fig. 3 D,E) compared to 3 hours (Fig. 3 A,B in B_ws_ state). At higher magnification, the average interaxial spacing of bundles is seen to be significantly larger in the B_ws_ phase (Fig. 3B) compared to bundles in the B_int_ phase (Fig. 3E). Microtubule pair distribution analysis of 6,603 MTs across several cross-sections and fields of view of TEM micrographs shows the difference in average interaxial spacing is 7.1 nm between the 3 hr (B_ws_) and 18 hr (B_int_) samples, comparable to differences in spacings measured through SAXS. Lastly, while the distance between neighboring MTs is heterogenous for both samples, there is less variation at 18 hours than at 3 hours. This is consistent with the observation seen in high magnification images (Fig. 3B and 3E) where the number of MT-MT bonds per MT is distinctly higher (i.e., MTs have a larger number of neighbors) in the B_int_ state than in the B_ws_ state. This also provides an explanation for the narrower peak widths (i.e. larger coherent domain sizes) observed via SAXS in the B_int_ phase.

**Figure 3.**
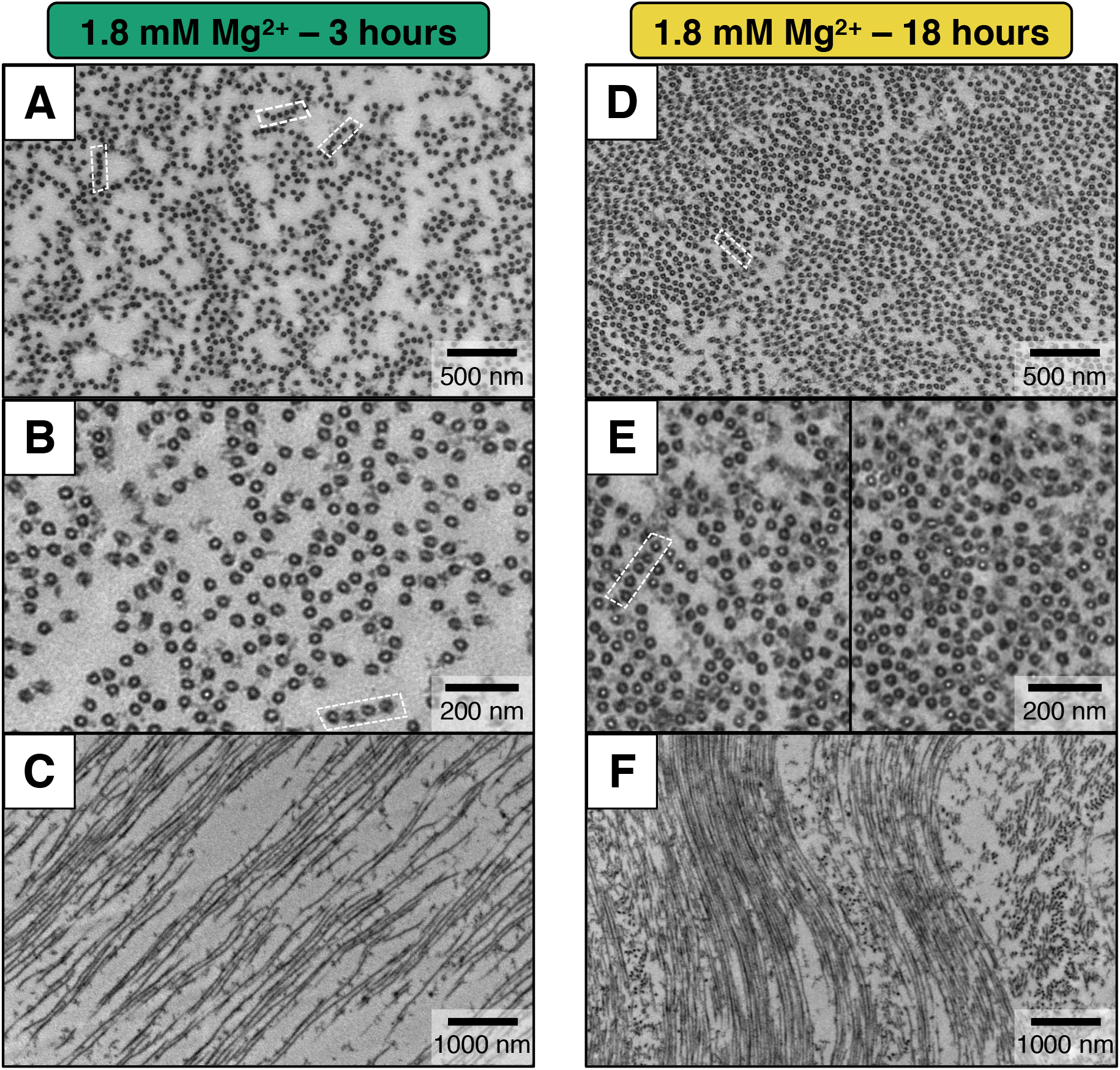
Plastic-embedded TEM confirms the existence of the B_ws_ and B_int_ bundled phases. Electron microscopy of microtubule assemblies prepared at 37 °C with mixtures of tau, tubulin, and 2mM GTP in standard buffer with 1.8 mM added Mg^2+^ and fixed after 3 hours (A-C) and 18 hours (D-E). Time points for sample fixation were selected based on SAXS data for samples prepared with identical conditions, where the wide-spacing B_ws_ and the intermediate B_int_ bundled phases were present at 3 and 18 hours, respectively. **(A**,**D)** Top panels depicting cross-sections at low magnification show the extent of MT bundling for both phases (B_ws_ state in A and B_int_ state in B). The images further show the propensity for MT bundles to arrange in linear arrays (dashed white boxes). **(B**,**E)** Higher magnification cross-sections at 3-hour and 18-hour timepoints highlight the larger MT-MT spacing in the B_ws_ compared to B_int_ phase and the larger number of MT-MT bonds per MT (and larger average number of neighbors for each MT) in the B_int_ compared to the B_ws_ phase. **(C**,**F)** Low magnification side views at 3-hour and 18-hour timepoints show that the width of the MT bundles (i.e. bundle size) is much larger in the B_int_ compared to the B_ws_ phase, and the spacing between MTs is smaller in the B_int_ compared to the B_ws_ phase, consistent with trends observed in SAXS data.

**Figure 4.**
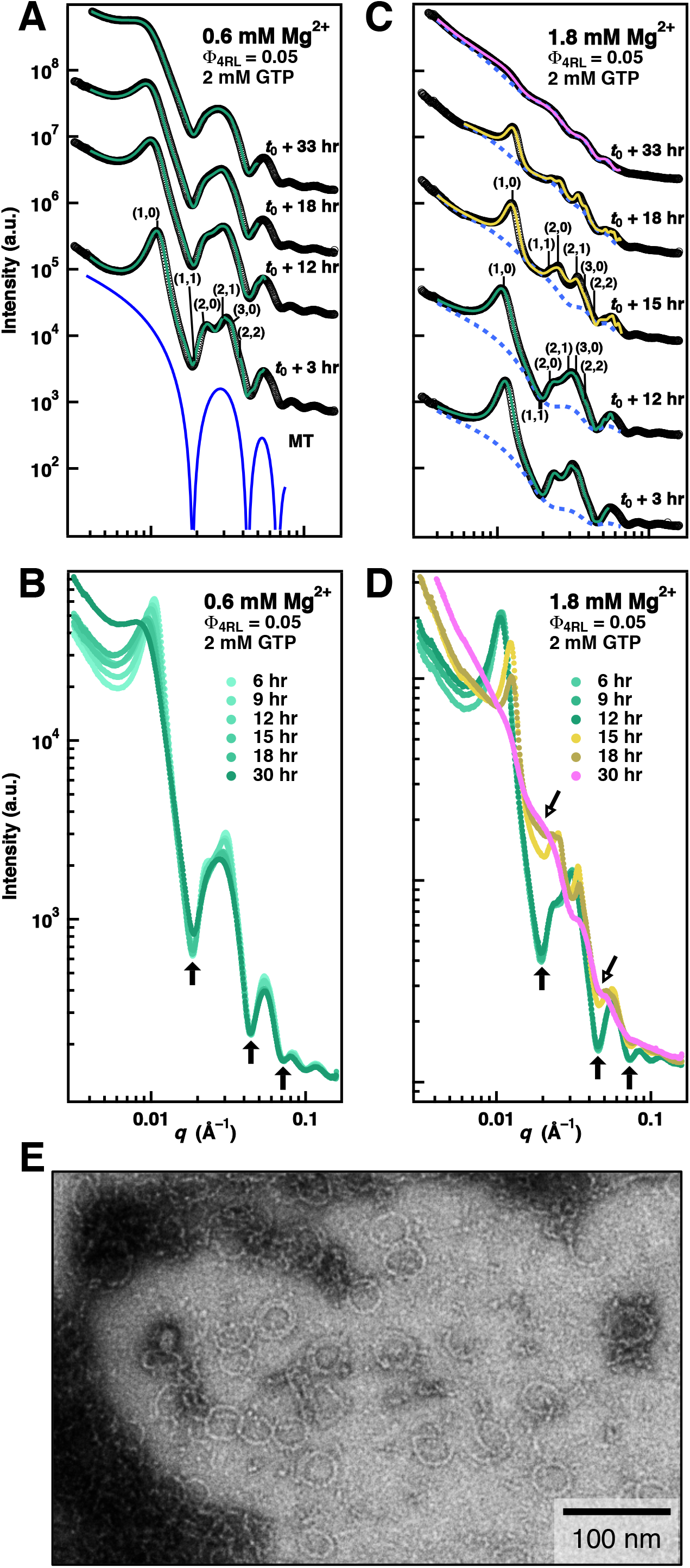
Time-dependent synchrotron SAXS data at 0.6 and 1.8 mM Mg^2+^ in the wide-spacing (B_ws_) and intermediate (B_int_) microtubule (MT) bundle states reveal prevalence of rings in the B_int_ state. (**A**) SAXS data (open circles, profiles offset for clarity) and corresponding fits (solid lines) of a sample at 0.6 mM Mg^2+^ (below c_lower_ ≈1.6 mM Mg^2+^) in the B_ws_ state (2D Hexagonal peaks indexed for profile at *t*_*0*_ +3 hrs). Bottom profile (solid blue curve) depicts the theoretical form factor of a single MT. (**B**) Data from (A) plotted without offset. Solid arrows point to minima in the MT form factor, highlighting nominal change in non-MT scattering over time. SAXS data (open circles, profiles offset for clarity) and corresponding fits (solid lines) at 1.8 mM Mg^2+^ (between c_lower_ ≈1.6 mM Mg^2+^ and c_upper_ ≈ 2.4 mM Mg^2+^) show a transition with increasing time from the B_ws_ (profiles at *t*_*0*_ +3 hrs and *t*_*0*_ +12 hrs) to the B_int_ (profiles at *t*_*0*_ +15 hrs and *t*_*0*_ +18 hrs) state and finally to the tubulin ring state (profile at *t*_*0*_ +33 hrs). Dashed lines are the non-MT scattering contribution obtained from fits as described in SI Appendix, SI Note S3. 2D Hexagonal peaks indexed for profiles at *t*_*0*_ +12 hrs (B_ws_ state) and *t*_*0*_ +15 hrs (B_int_ state). (**D**) Data from C without offset. An abrupt increase in tubulin ring scattering fills in the minima of the MT Form Factor (open arrows) at the time of transition between *t*_*0*_ +12 and *t*_*0*_ +15 hours. The fit lines in A-D are color-coded (green, yellow, and magenta represent the B_ws_, B_int_, and tubulin ring states, respectively). (**E**) Parallel whole-mount TEM image taken at t_0_ + 18 hrs of a sample prepared with 1.8 mM Mg2+ added, showing abundance of single tubulin rings.

TEM experiments additionally reveal structural features on a larger length scale beyond the resolution of SAXS. Specifically, low-magnification large field-of-view slices on the micron-scale along the MT length (Fig. 3C and F) show that MT bundles form notably larger extended arrays in the bundled intermediate state (Fig. 3F, 18 hrs) compared to the wide-spacing state (Fig. 3C, 3 hrs).

In agreement with plastic-embedded TEM measurements of Chung et al. (*35*) (with no added Mg^2+^ to the buffer), few well-defined hexagonal arrays, especially in the wide-spacing state, are observed. Instead, there is an apparent preference for MTs to form linear, string-like bundles (Fig. 3 A,B,D,E white boxes) reminiscent of fascicles found within the axon-initial-segment (SI Appendix, Fig. S1). Even within regions of high microtubule density (Fig. 3D), where the probability of MT-MT interactions is higher, stacks of linear arrays or branched chains of MTs continue to dominate over true hexagonal bundles. It is important to note that the fixing, staining, and plastic embedding process for TEM does distort the lattice between MTs to some extent. Nonetheless, this observed preference for 1-dimensional bundling between MTs despite close lateral proximity of the linear arrays may be due to breaking of cylindrical symmetry with a non-uniform distribution of tau on the MT surface. This is consistent with reports that tau forms phase separated dimeric/trimeric complexes bound to the surface of MTs (*42-44*).

### SAXS line-shape modeling of the B_ws_ and B_int_ microtubule bundled phases and tubulin rings

To quantify the observed divalent cation-mediated spike in MT depolymerization in SAXS at the B_ws_ to B_int_ transition (i.e. filling in of profile’s local minima, Fig. 2A,C), scattering data was fit to a model profile, *I*(q) (Eq. 1), which consisted of three separate terms modeling hexagonally bundled microtubules (*35, 41, 45*), tubulin rings, and background scattering(*46*) to account for a constant background level and scattering from unpolymerized tubulin oligomers (i.e. oligomers not in the MT lattice) with average curvature much less than that of the closed rings (SI Appendix, SI Note S3):

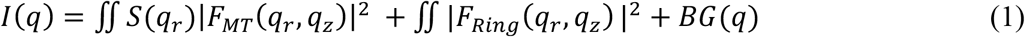

The first term in *I*(q) consists of the structure factor S(q_r_) of the bundled MT lattice multiplied by the form factor of a MT (|F_MT_(q_z_,q_r_)|^2^) and is averaged over all orientations in q-space (q_r_, q_z_ are wavevectors perpendicular and parallel to the MT cylinder axis). S(q_r_) (SI Appendix, SI Note S3, Eq. (2)) was modelled as a sum of square Lorentzian functions (*45*) at each reciprocal lattice vector q_*hk*_ = q_*10*_(h^2^+k^2^+hk)^1/2^ with amplitude A_*hk*_ and peak width κ_*hk*_. The form factor (SI Appendix, SI Note S3, Eq. (3)) was calculated by modeling the MT as a hollow cylinder with uniform electron density(*33, 47*) with wall width set to *w* = 49 Å (*48*), and length L_MT_ fixed at 20,000 Å (larger than the resolution of our wavevector) (*47*). The MT cylinder’s inner radius, *r*_in_, was the only fit parameter in the form factor. The second term in *I*(q) which accounts for the scattering contribution from uncorrelated tubulin rings (SI Appendix, SI Note S3, Eq. (4)) was also modelled as the Fourier transform of a hollow cylinder but of length L_Ring_ = 49 Å (width of one tubulin monomer) with the ring’s inner radius and scattering amplitude (A_ring_) being fit parameters.

It is important to note that since the first order Bessel functions of the MT form factor (*J*_1_ terms in Eq. (3), SI Appendix, SI Note S3) equal 0 for distinct values of *q*_r_ (given *r*_in_), the scattering contribution from MTs (both bundled and unbundled) to the total raw scattering profile will also approach zero at these points, resulting in the deep local minima observed in the theoretical scattering from a single MT (Fig. 4A, bottom blue curve). Therefore, since scattering from all tubulin within the MT lattice is suppressed at these distinct values of *q*_r_, the measured intensity at these points within the raw data must be almost entirely due to the scattering from unpolymerized tubulin oligomers plus a q-independent background. Furthermore, any observed changes in the scattering intensity at these minima over time must, therefore, be due to changes in unpolymerized tubulin content.

### Distinct types of unpolymerized tubulin oligomers in the B_ws_ and B_int_ bundled phases

Fits to the time-dependent SAXS data of reaction mixtures polymerized with 0.6 and 1.8 mM added Mg^2+^ (Fig. 4, A–D) highlights the differences in scattering from unpolymerized tubulin oligomers between the B_ws_ and B_int_ states for Mg^2+^ samples (similar to behavior found for Ca^2+^ samples, Fig. 2 A, C). The solid lines through the SAXS profiles in Fig. 4(A,C) are fits of the data to *I*(q) (Eq. 1). For 0.6 mM Mg^2+^ (below c_lower_ ≈1.6 mM Mg^2+^) the B_ws_ state is stable for the duration of the experiment (Fig. 4A) despite indications of gradual MT depolymerization due to ongoing partially suppressed dynamic instability. Overlapping the SAXS profiles without offset shows that there is a slight increase in scattering intensity at the first local minimum over time (Fig. 4B, solid arrows), but there is a lack of increased scattering intensity at all other minima. This scattering change is due to an increase in MT depolymerization products over time. However, these tubulin products cannot be in the ring conformation because tubulin ring scattering (|F_Ring_|^2^, SI Appendix, SI Note S3, Eq. (4)) would result in a linear increase of all local minima. Late time points (up to 72 hours) in control samples with no added divalent cation show this phenomenon most clearly, as the intermediate B_int_ state is not observed and scattering from tubulin rings is absent despite significant MT depolymerization (SI Appendix, Fig. S2). Thus, for data in the B_ws_ state (Fig. 4A), the ring scattering amplitude, A_ring_, is zero, and *I*(q) that fits the data well is due to the MT bundle scattering plus background scattering from lower curvature tubulin oligomers (contained in the third term in Eq. (1)).

For Mg^2+^ concentrations between c_lower_ and c_upper_ (≈1.6 to ≈2.4 mM Mg^2+^), the B_ws_ state is stable for many hours with characteristically little tubulin ring formation (Fig. 4, C (*t*_0_ +3 hrs and *t*_0_ +12 hrs) and D without offset in SAXS data (*t*_0_ +6 hrs, *t*_0_ +9 hrs, and *t*_0_ +12 hrs)). However, transitioning to the intermediate bundled state (Fig. 4C, *t*_0_ +15 hrs and *t*_0_ +18 hrs, and Fig. 4D, *t*_0_ +15 hrs, *t*_0_ +18 hrs, and *t*_0_ +30 hrs) coincides with a sudden spike in scattering intensity at all form factor minima, indicated by solid arrows in Fig. 4D. This distinct change in scattering (between 12 and 15 hours for 1.8 mM Mg^2+^), unlike that of the B_ws_ state, is well described by an increase in scattering from tubulin rings (Fig. 4C, blue dashed lines). Plots of the raw scattering data without offset in Fig. 4D for 1.8 mM Mg^2+^ show the remarkable overlap around the local minima of profiles at the *t*_0_ +18 and *t*_0_ +30 hrs time points, where oscillations in SAXS at *t*_0_ +30 hrs are well described by the theoretical scattering profile of a tubulin ring with *r*_in_ = 16.3 nm. Whole-mount TEM images confirm observations made with SAXS, showing a relative abundance of individual and aggregated tubulin rings at 18 hours in the B_int_ state with 1.8 mM Mg^2+^ (Fig. 4E).

Together the data indicates that changes to the bundled MT lattice only manifest upon the observed proliferation of tubulin rings. The fitted data from all samples observed in the ring state, with a typical example shown in Fig. 4C at *t*_0_ +33 hrs, are consistent with a single ring model with an average inner ring radius of *r*_in_ of ≈16.3 nm (ranging between 15.7 and 17.4 nm). Modeling the reported double ring state did not produce convincing fits and resulted in *r*_in_ values far from those observed in prior TEM work (*49*). Whole mount TEM of rings in the intermediate state is consistent with this SAXS analysis, where double rings can be observed but the vast majority are single tubulin rings (Fig. 4E).

### Abrupt proliferation of tubulin rings occurs at the same time as the decrease in microtubule wall-to-wall spacing, signaling the B_ws_ to B_int_ transition

Figure 5 summarizes the results of fitting parameters obtained from fits of the SAXS data to *I*(q). Comparing the MT wall-to-wall distance *d*_w-w_ to the amplitude of the ring scattering (A_ring_) of samples at different Ca^2+^ and Mg^2+^ concentrations as a function of time, highlights the striking synchronization between the proliferation of tubulin rings and abrupt drop in *d*_w-w_ upon transitioning from the B_ws_ to the B_int_ state (Fig. 5, compare time points of arrows in A to those in B, and arrows in C to those in D).

**Figure 5.**
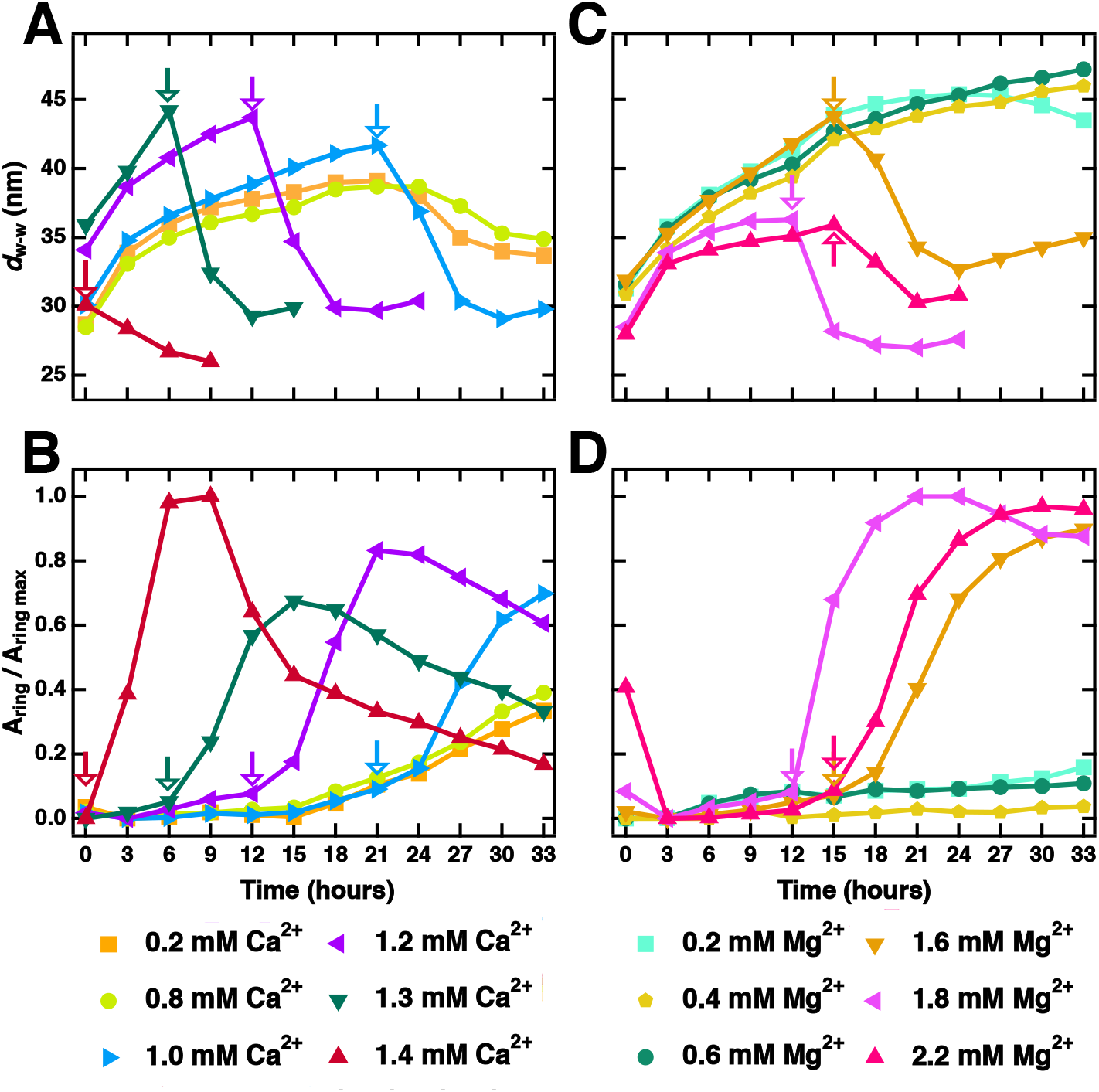
Change in the microtubule (MT) wall-to-wall spacing, upon transitioning from the wide-spacing (B_ws_) to the intermediate (B_int_) MT bundle state, correlates with tubulin ring proliferation. (**A** and **C**) Wall-to-wall spacings (*d*_w-w_ = a_h_ – 2[*r*_in_ + *w*]) plotted as a function of time for the series of Ca^2+^ (A) and Mg^2+^ (C) samples (SAXS data shown in Figures 2 and 4, respectively). Some data points omitted for clarity. Arrows indicate the latest time point at which the B_ws_ state is observed before the sample transitions to the B_int_ state. (**B** and **D**) Fitted tubulin ring scattering amplitude plotted as a function of time for the same series of Ca^2+^ (B) and Mg^2+^ samples. Amplitudes are normalized by the maximum ring scattering measured. Arrows indicate the latest time point at which the bundled wide-spacing (B_ws_) state is observed before the sample transitions to the bundled intermediate (B_int_) state, with increases in the tubulin ring scattering amplitude observed after. Comparisons between (A) and (B) for the Ca^2+^ series and between (C) and (D) for the Mg^2+^ series show that the onset of tubulin ring proliferation occurs concurrently with the transition from the B_ws_ to the B_int_ state. Time “0” on the x-axis corresponds to t_0_, as defined in Fig. 2.

For samples in the B_ws_ state, *d*_w-w_ increased rapidly at early time points due to a relaxation from sample centrifugation (Sample Preparation in Methods) to spacings ≈45 nm (Fig. 5A,C) and more slowly thereafter, reaching values up to *d*_w-w_ ≈49 nm at 33 hours. Unlike in the B_ws_ state, *d*_w-w_ spacings are comparatively stable over time after transitioning fully into the B_int_ state (i.e. starting with time points between 3 and 6 hrs after arrows in Fig. 5A,C). The average stabilized wall-to-wall distance for B_int_ samples that transitioned into the intermediate state hours after preparation was 29.8 nm and was independent of *d*_w-w_ in the B_ws_ state at the timepoint directly preceding the transition (shown with arrows in Fig. 5A,C). Similarly, samples initially observed in the intermediate B_int_ state at *t*_0_ (see Fig 2D, yellow region at time = 0) show no relaxation or increase in *d*_w-w_ over time (unlike what is found in the B_ws_ state), instead converging to average spacing *d*_w-w_ = 30.4 nm for Mg^2+^ and 26.4 nm for Ca^2+^. The stability of *d*_w-w_ values measured in the B_int_ state relative to those in the B_ws_ state (which may fluctuate with time) implies that stronger interactions dictate MT-spacing in the B_int_ compared to the B_ws_ state. This implication is further substantiated as not only does the *d*_w-w_ spacing drop concurrently with the initiation of tubulin ring formation (i.e., time points within the 3 hrs after arrows in Fig. 5A, C), but also the coherent domain size in the B_int_ state grows significantly larger over the same time period, with average growth of ≈41 percent in the 6 hours following the transition (SI Appendix, Fig. S3).

Significantly, SAXS data shows that the B_int_ state (Fig 2D, yellow region) consists of a two-phase system where MT bundles (first phase) co-exist with a steady influx of tubulin rings (second phase, which eventually dominates) due to ongoing MT depolymerization of a fraction of MTs (either isolated MTs or MTs at the periphery of bundles where fewer cross-bridges to neighboring MTs exist). The drop in *d*_w-w_ together with the increase in MT bundle domain size, precisely when increasing amounts of rings and curved tubulin oligomers are being produced, suggests that tubulin oligomers directly affect the bundling of MTs and drive the B_ws_ to B_int_ transition observed through SAXS and TEM.

### Enhancement or suppression of unpolymerized tubulin oligomers dictates transitions between the wide-spacing (B_ws_) and intermediate (B_int_) bundled microtubule states

To directly test the effect of enhanced unpolymerized tubulin oligomers on MT bundling, time-dependent SAXS experiments were devised to modulate dynamic instability. Protofilaments (tubulin oligomers) in excess GTP are known to adopt a higher curvature conformation at low temperature (*7, 50*). Therefore, lowering the temperature of the sample environment below a critical temperature for polymerization pushes the tubulin dynamic equilibrium toward depolymerization, effectively increasing the tubulin oligomer content in solution. Monitoring a control sample (i.e., with no added divalent cations) in the B_ws_ state while the temperature was quickly dropped from 37 °C to 5 °C revealed that the wide-spacing state (which consists of MT bundles co-existing with tubulin oligomers from the ongoing suppressed dynamic instability) was stable down to temperature readings of 19 °C with no increase in scattering from depolymerized tubulin (Fig. 6A, bottom two profiles). However, below 19 °C, signs of the B_int_ state abruptly appear, including the thematic shift in hexagonal Bragg peaks to higher *q* coinciding with a sudden increases in tubulin scattering at 13.8 °C and at 5.8 °C (evidenced as the filling in of the minima in the profiles in Fig. 6A at these temperatures compared to profiles at 34.5 °C and 19.0 °C). In the B_int_ state observed at these lower temperatures (Fig. 6A, 13.8 °C and 5.8 °C), *d*_w-w_ decreased to a minimum value of 26.1 nm (change in wall-to-wall distance, Δ*d*_w-w_ = 8.8 nm). Scattering from the tubulin ring state was dominant at temperatures below 5.8 °C (Fig. 6A 4.9 °C). This result further demonstrates the correlation between increased free tubulin oligomer content and MT bundle rearrangement as the system transitions from the B_ws_ to the B_int_ state. Additionally, it shows that *d*_w-w_ can be altered independent of any ionic changes to the buffer and only requires the presence of different concentrations of free tubulin. In contrast, for the previously described experiments performed at 37°C at 2mM GTP, the increase in curved tubulin oligomers was induced by increasing the divalent cation concentration above a critical concentration, which precipitated transitions from the B_ws_ to the B_int_ and tubulin ring states.

**Figure 6.**
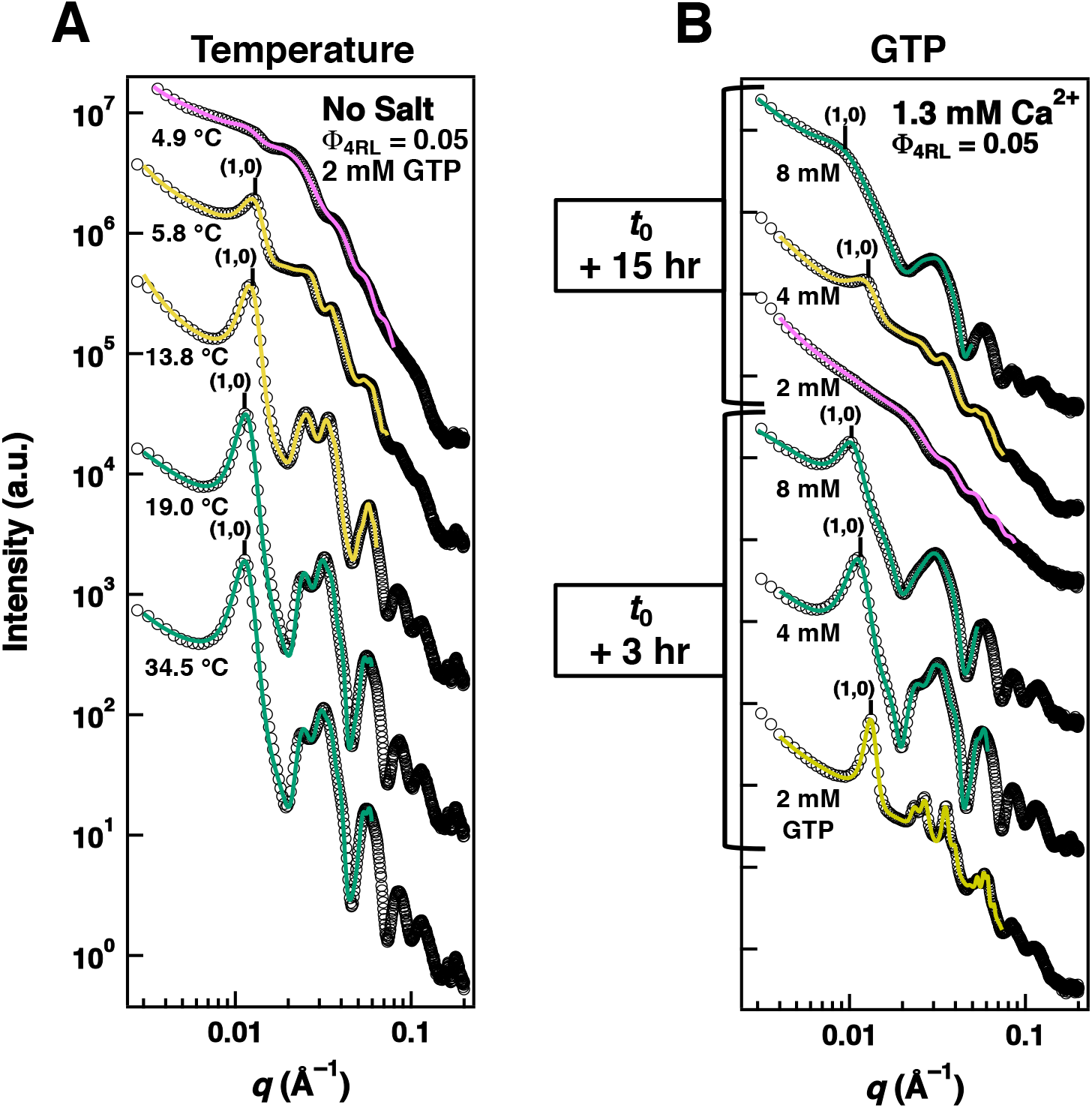
Synchrotron SAXS data reveal the stability of the bundled wide-spacing state depends on temperature and concentration of GTP. (**A**) SAXS data (open circles) and corresponding fits (solid lines) for a control sample prepared with no added divalent cations while the temperature of the sample holder enclosure was rapidly dropped from 37 °C to below 5 °C. Temperatures were recorded simultaneously with each SAXS exposure and are displayed next to each scattering profile. Profiles represent the first and last time point at which the bundled wide-spacing (34.5 and 19.0 °C, respectively), the bundled intermediate (13.8 °C and 5.8 °C, respectively), and the tubulin ring state (4.9 °C) were observed. (**B**) SAXS data (open cricles) and corresponding fits (solid lines) at *t*_*0*_ + 3 and *t*_*0*_ + 15-hour time points for samples at 37°C prepared with 1.3 mM CaCl_2_ and either 2, 4, or 8 mM GTP. The data show that increasing GTP concentrations favors the B_ws_ over the B_int_ state at both time points, and the B_int_ over the tubulin ring state at time point *t*_*0*_ + 15-hour. The fit lines in both A and B are color-coded (green, yellow, and magenta represent the B_ws_, B_int_, and tubulin ring states, respectively).

To separately test the effect of decreased tubulin oligomer content, reaction mixtures were prepared at 37 °C with 1.3 mM Ca^2+^ and either 2, 4, or 8 mM GTP, then monitored via SAXS for 33 hours. These samples reproduced the time-dependent trends of the kinetic phase diagram in Fig. 2D; however, increasing GTP, which is known to suppress the formation of unpolymerized tubulin oligomers (by incorporation of GTP-PFs in the MT lattice), increased the stability of the wide-spacing state over the intermediate state and the intermediate state over the tubulin ring state. As seen in Fig. 6B, samples prepared with 4mM GTP did not transition from the B_ws_ to the B_int_ state by t_0_ +3 hrs and did not transition from the B_int_ to the ring state by t_0_ +15 hrs (unlike samples prepared with 2mM GTP). Furthermore, samples prepared with 8 mM GTP remained in the wide-spacing B_ws_ state for the duration of the experiment. Taken together, these results show an unequivocal dependence of MT bundling characteristics (in particular, the change in the wall-to-wall spacings in the B_ws_ and B_int_ states) on free tubulin content.

### Complexes of tubulin oligomers and tau act to cross-bridge microtubules and stabilize microtubule bundles

Plastic embedded TEM side-views along the MT length provide potential insight to the structural components cross-bridging neighboring MTs and reveal a possible explanation for the B_int_ phase observed through SAXS. Close inspection of bundles in the B_ws_ phase at 3 hours at lower (Fig. 7 A,B) and higher (Fig. 7 C-E) magnifications reveals an extended intervening network of crosslinked proteins connecting MTs (Fig. 7 C-E). Remarkably, cross-bridges are seen not only connecting MTs within bundles (Fig 7 C-E, solid black arrows), but also connecting neighboring bundles to one another (Fig 7 C-E, dashed black arrows). Evidence of these tethers are also seen within the high magnification cross-sectional views shown in Fig. 3, with more crosslinks present in the B_int_ state (Fig. 3E) compared to the B_ws_ state (Fig. 3B).

**Figure 7.**
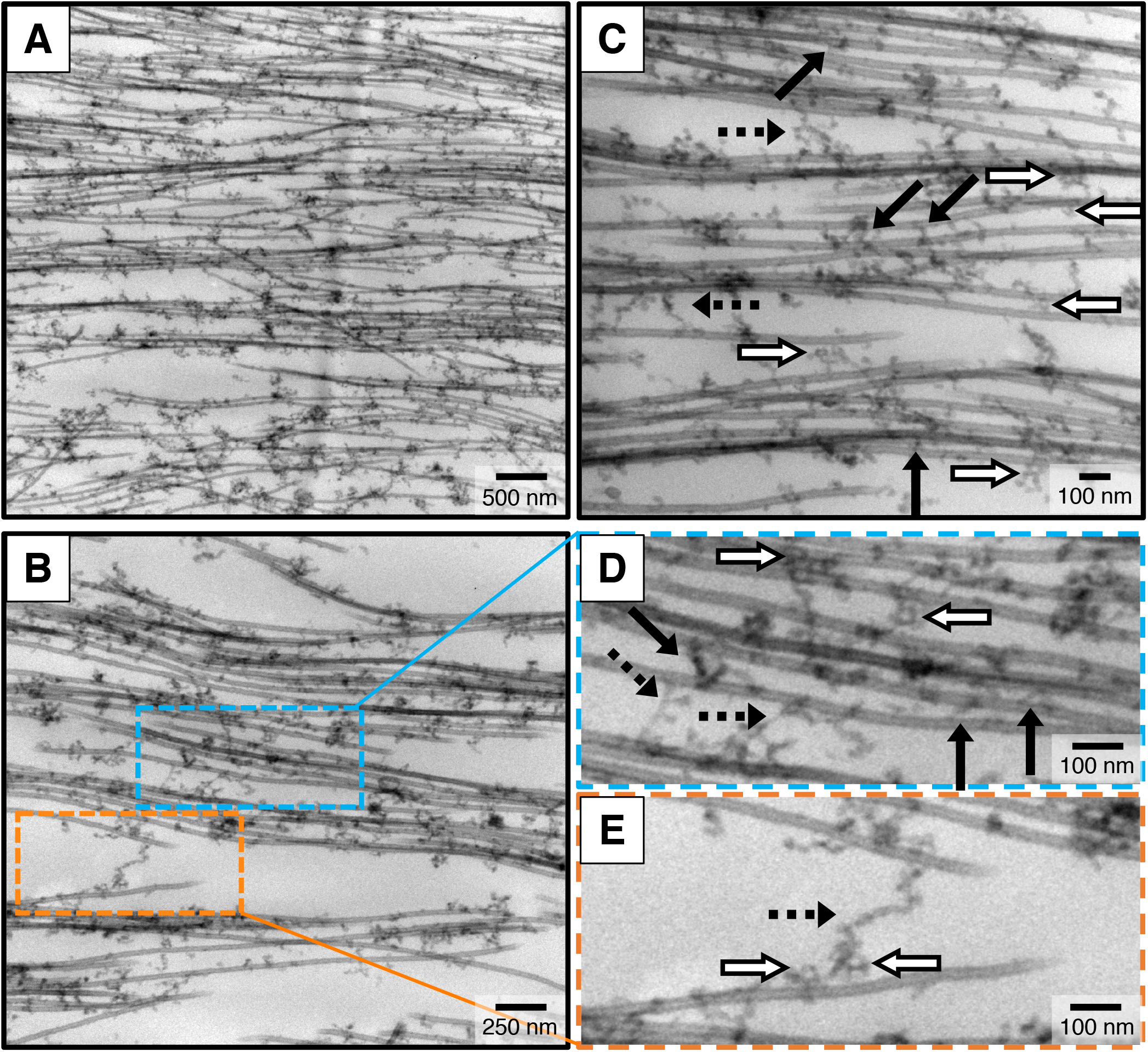
Plastic-embedded TEM provides evidence that complexes of tubulin oligomers and tau cross-bridge bundled microtubules. Electron microscopy with increasing magnification of microtubule assemblies prepared from mixtures of tau, tubulin, and 2 mM GTP in standard buffer with 1.8 mM added Mg^2+^ at 37 °C and fixed after 3 hours. **(A**,**B)** At low and intermediate magnification, TEM images provide evidence that a network of filamentous proteins between bundled MTs is the linking medium that stabilizes MT bundles. **(C-E)** At higher magnifications the filamentous proteins of the network are seen to form MT-MT cross-bridges both within (black arrows) and between (black dashed arrows) bundled domains. The morphology of these cross-bridges is consistent with ≈5 nm wide semi-flexible tubulin oligomers. Tubulin ring structures (white arrows) are also present within the protein network. (D, E are expanded views of sections in B with blue and orange outlines.) Complexes of tubulin oligomers and tau are also observed coating MTs but not forming MT-MT cross-bridges.

While cross-bridges between bundled MTs in cells have been previously reported but attributed only to tau (*8, 9, 51*), intrinsically disordered proteins such as tau (≈0.5 nm in width) are too thin to be visualized by TEM in our plastic embedded preparations, implying that tau alone cannot make up the MT cross-bridge. In contrast, the morphology and dimension of the cross-bridges observed in TEM are consistent with tubulin oligomers, existing both as semi-flexible filaments (black and black dashed arrows in Fig. 7 C-E) and as tubulin ring structures (white arrows in Fig. 7 C-E). Thus, TEM data provides direct evidence that tubulin oligomers (which may include tubulin rings) are a central component of the observed intervening network between MTs and suggests that tau’s role in MT bundling is to act as the “glue” (through the binding repeats) which connects tubulin oligomers. Also seen in TEM are complexes of tubulin oligomer and tau bound to isolated MTs not in bundles.

## Discussion

In summary, our combined SAXS and TEM data lead us to propose a model where complexes of tubulin oligomers and tau form an intervening network that cross-bridge MTs into bundles. We propose that bundling occurs due to *coded-assembly* where tau’s MT binding repeats link together αβ-tubulin oligomers in the intervening network and also αβ-tubulin oligomers near the MT surface to αβ-tubulin in the MT lattice (Fig. 1B). The model presents an important revision to current dogma where cross-bridging between MTs is attributed entirely to tau (*8, 9, 12, 35, 51-53*).

Our model is consistent with all of the observed characteristic of the GTP-stabilized B_ws_ and B_int_ MT bundled states, and the transition between them, observed in time-dependent SAXS. The abrupt influx of (tau coated) tubulin rings and curved tubulin oligomers (due to Mg^2+^ or Ca^2+^ mediated MT depolymerization of a fraction of MTs in the bundled state) drives the transition from B_ws_ to B_int_ by increasing the average number of tubulin-tau cross-bridges (seen in TEM) in remaining transiently stable B_int_ bundles. The enhanced cross-bridging simultaneously decreases *d*_w-w_ while increasing the bundle domain size in the B_int_ state seen in SAXS.

The model reconciles years of contradicting reports regarding tau’s role in bundling. While numerous early publications pointed to MT bundling as one of the many roles of tau in cells (*8, 9, 12, 51*), more recent studies reported that the use of the MT-stabilizing drug paclitaxel, which severely reduces free tubulin oligomers (at paclitaxel/tubulin-dimer molar ratios of Λ_paclitaxel_=1/1), suppresses MT bundling by all six tau isoforms (*47*). Furthermore, follow-up SAXS and TEM experiments showed that reducing paclitaxel to ratios below Λ_paclitaxel_ ≈1/8, restores free tubulin and restores MT bundles in the presence of tau (*54*). These findings are consistent with our central discovery; namely, that bundling of MTs with tau requires free tubulin oligomers.

In contrast, tau-only models of MT bundling would require a highly extended conformation for the projection domain (PD) of tau (Fig. 1A). In one model, the bundling is associated with short-range attractions between cationic and anionic residues of weakly penetrating tau PDs near the midplane layer between opposing microtubule surfaces (*35*). For the SAXS data presented in this paper, the physical diameter of the projection domain of 4RL-tau D_Phys_ = 2R_Phys_ = 2(5/3)^1/2^R_g_ ≈9.6 nm (*55*). (Here, the radius of gyration R_g_ ≈3.8 nm as measured in solution SAXS (*56, 57*)). This would lead to a MT wall to wall spacing *d*_w-w_ ≈19.2 nm for two opposing (weakly overlapping) tau chains on neighboring MTs. Thus, this tau-only model would require the tau’s PD to be extended by about a factor of two in order to account for the average *d*_w-w_ ≈38 nm observed in SAXS. In another tau-alone electrostatic-zipper model based on the dipolar nature of 4RL-tau (*53*), the cross-linking of two MTs is achieved by the complete overlap of anti-parallel Tau PDs. In this latter model, the PD would have to be stretched by about a factor of four to account for *d*_w-w_ (approaching the contour length of 4RL-Tau’s PD ≈54 nm with 150 amino-acid residues).

Extending the PD by factors of 2 or 4 is inconsistent with traditional polyelectrolyte theories (*58*) for chain stretching in the mushroom regime (at low tau/tubulin dimer molar ratio), and would require *sequence-specific* theories of highly stretched polypeptide chains at physiological salt concentrations (≈150mM 1:1 salt). SAXS measurements of tau-coated paclitaxel-stabilized MTs at Λ_paclitaxel_ = 1/1, which do not bundle due to the lack of sufficient free tubulin and require an applied osmotic pressure in order to bundle (*33*), exhibit *d*_w-w_ spacings in the range of ≈3.5-4nm close to the radius of gyration of tau’s PD and consistent with polyelectrolyte theory. In contrast, similar osmotic pressure studies of MT bundles in the B_ws_ state with available free tubulin oligomers (*35, 41, 45*) show that the wide spacing between MTs is retained even at high pressures consistent with our model where tubulin oligomers are components of the MT cross-bridges.

In our model, the wall-to-wall spacing *d*_w-w_ is largely set by the average radius of curvature of tau-coated curved tubulin oligomers and tubulin rings (outer diameter ≈40 nm). Furthermore, because the binding interactions between tau and tubulin oligomers are specifically encoded by the MT binding repeats of tau, one expects MT bundles to be only weakly dependent on added monovalent salts. Indeed, *d*_w-w_ is found to be essentially constant over a wide range of salt concentrations up to ≈150 mM KCl added to the buffer, which already contains ≈100 mM of a 1:1 salt (SI Appendix, Fig. S4, left). Additionally, in SAXS experiments designed to test the effects of tau coverage on MT wall-to-wall spacing, we find that *d*_w-w_ shows little change in spacing between samples, despite tau-tubulin dimer molar ratios ranging from Φ_4RL_ = 1/100 (in the mushroom regime) to Φ_4RL_ = 1/5 (in the brush regime) (SI Appendix, Fig. S4, right panels). This is consistent with our model but not consistent with a tau-alone model where one would expect an increase in *d*_w-w_ upon surpassing the mushroom-to-brush threshold Φ_4RL_ ≈1/10 (*33*). The encoded binding interaction also better accounts for MT’s ability to bundle at tau tubulin ratios as low as Φ_4RL_ = 1/100.

Together our results draw attention to the essential role of unpolymerized tubulin on MT bundling in a minimal dissipative reaction mixture at 37°C containing tubulin, tau, and GTP and highlight the necessity of operating within physiologically relevant experimental conditions as even minor differences in buffer conditions or temperature may have significant repercussions when drawing conclusions on tau’s bundling function. Our finding that divalent cations, near average physiological Mg^2+^ concentrations, can destabilize dynamic, steady-state MT bundles at 37 °C, despite the presence of tau and excess GTP, is unexpected. In particular, at concentrations above a critical divalent concentration c_lower_ (≈1.6 mM Mg^2+^), MT bundles undergoing suppressed dynamic instability abruptly become unstable towards the formation of tau-coated tubulin rings, effectively halting dynamic instability in the reaction mixtures. This suggests a mechanism where MT growth and stability can be heavily modulated within cells through slight fluctuations in local divalent cation concentrations, with potential consequences for cargo transport in axons. Additionally, we expect the value of c_lower_ to be modulated by disease-relevant post-translational modifications like tau hyperphosphorylation or cleavage.

We expect other members of the vertebrate family of MAPs with similar MT binding repeats in their C-terminal domains(*59*), including MAP2 in dendrites and MAP4 in non-neuronal cells, to behave in a manner similar to MAP tau with respect to interactions with MTs. For example, the spacing between MTs in bundles formed by MAP2 expressed in cells is wider than that with tau (*8, 9, 51, 52*). MAP2 has a significantly longer projection domain (with a correspondingly much larger negative electrostatic charge) compared to tau and the larger spacing may possibly be due to a lower average curvature of tubulin oligomers complexed with MAP2 in the intervening network. It would be important to address the precise nature of MT bundling due to MAP2 in cell-free studies of minimal reaction mixtures similar to what we have done for tau.

We should point out that previous studies show that, *in vivo*, the MT binding protein TRIM46 localizes to the proximal axon region promoting parallel MT bundles oriented with their plus end out (*60-62*). The reported function of TRIM46 would be synergistic to our finding in cell free reaction mixtures where tubulin-tau complexes cross-bridging MTs in bundles, in the absence of any other protein or molecular component, result in the linear string-like ordering of MTs found in the axon-initial-segment in mature neurons (compare Fig. 3A to Fig. S1, SI Appendix).

The implications of our discovery of an intervening network of complexes of tubulin oligomer and tau should spur further studies. For example, one major function for bundled MT fascicles at the axon-initial-segment (AIS) is as a filter for MT-based cargo trafficked between the soma and axon. An intervening tubulin-tau network stabilizing MT fascicles should lead to more efficient retrograde diffusion barrier properties in capturing and preventing tau leakage outside of the axon. In the event where tau is biochemically altered or fragmented (as happens in Alzheimer’s disease and other tauopathies) the barrier properties in the AIS could be modulated by alterations to the intervening network with significant implications for missorting of tau to the somatodendritic compartment and neurodegeneration. Finally, complexes of tubulin oligomers and tau of the intervening network, or those bound to MTs not in bundles, may represent an important site of chemical modification or fragmentation of tau by enzymes leading to aberrant tau behavior and nucleation and growth of tau fibrils in tauopathies.

## Methods

### Tubulin and Tau Purification

Tubulin was purified from bovine brain as described previously(*32*) (SI Appendix, SI Note S1.1). Tau was expressed in and purified from BL21(DE3) pLacI cells (Invitrogen), following standard procedures (*63*) (SI Appendix, SI Note S1.2).

### Sample Preparation

Purified tubulin was diluted into PEM50 and mixed on ice with PEM50 solutions containing GTP, tau, and additional salt as indicated. Final polymerization mixtures contained 3.5 – 4.0 mg/mL tubulin in 40 – 50 uL of buffer. Sample tubes containing polymerization mixtures were placed in a 37 °C water bath for 30 minutes for polymerization to reach dynamic equilibrium. The MT-tau samples were then prepared for experiments as follows. For SAXS, samples were directly loaded into 1.5-mm diameter quartz mark capillaries (Hilgenberg GmbH, Malsfeld, Germany) after polymerization. Capillaries were subsequently spun in a capillary rotor in a Universal 320R centrifuge (Hettich, Kirchlengern, Germany) at 9,500 x *g* and 37 °C for 30 minutes to form protein-dense pellets suitable for SAXS. Pelleted capillaries were then held at 37 °C in a custom-made temperature-controlled sample holder for data acquisition. Sample preparations for whole-mount and plastic embedded TEM were described previously (*35, 41, 45*) with minor changes (SI Appendix, S2 Note S2).

### X-Ray Scattering and Analysis

SAXS experiments were performed at beamline 4-2 of the Stanford Synchrotron Radiation Lightsource at 9 keV using a custom-made temperature-controlled sample holder. Experiments were performed at 37 °C unless otherwise noted. For the temperature ramp down experiment, initial data was taken at 37 °C. Then the temperature controller was set to 5 °C, which was reached over the course of <7 minutes. A needle temperature probe was inserted into a water-filled quartz capillary placed in the sample holder for instantaneous temperature recordings during the ramp down. Quantitative line-shape analysis was performed (SI Appendix, SI Note S3).

### Transmission Electron Microscopy

All data were taken at 80 kV using the JEM 1230 (JEOL) Transmission Electron Microscope at the University of California, Santa Barbara.

## Acknowledgements

This work was supported in part by the US National Science Foundation, Division of Materials Research, under award DMR-1807327 (CRS, phase behavior in biomaterials) and the Department of Energy, Office of Basic Energy Sciences, Division of Materials Sciences and Engineering, under award DE-FG02-06ER46314 (CRS and YL, charged bio-assemblies inspired by nature). PJC acknowledges support from the US National Science Foundation under award DMR-2104854. SCF acknowledges support from the US National Institutes of Health grant NS-35010 and the Academic Senate of the University of California at Santa Barbara. MCC acknowledges support from the National Research Foundation of Korea NRF-2018R1A2B3001690, and interactions with *KAI-NEET* Institute (affiliated to KAIST), and the Asian Pacific Center for Theoretical Physics (APCTP).

## Author contributions

C.R.S., C.S., B.F., P.K., and Y.L. designed research; H.P.M. purified tubulin; C.S., B.F., C.T., and R.L.B. provided tau protein isoforms from plasmid preparations; C.S., B.F., P.K., and X.G.A. performed research; P.K. analyzed SAXS data; C.R.S., P.K., B.F., and Y.L. modeled data; P.K., B.F., and C.R.S. wrote the paper; Y.L., P.J.C., R.L.B., S.C.F., M.C.C., L.W., C.T., and H.P.M. provided additional writing input; and Y.L., S.C.F., M.C.C., and L.W. provided critical suggestions on presentation of X-ray and TEM microscopy data.

**The authors declare no competing interest**.

## References and Notes

1. Desai, T. J. Mitchison, Microtubule polymerization dynamics. Annu Rev Cell Dev Biol 13, 83–117 (1997).

2. T. Muller-Reichert, D. Chretien, F. Severin, A. A. Hyman, Structural changes at microtubule ends accompanying GTP hydrolysis: information from a slowly hydrolyzable analogue of GTP, guanylyl (alpha,beta)methylenediphosphonate. Proc Natl Acad Sci U S A 95, 3661–3666 (1998).

3. G. M. Alushin et al., High-resolution microtubule structures reveal the structural transitions in alphabeta-tubulin upon GTP hydrolysis. Cell 157, 1117–1129 (2014).

4. A. A. Hyman, D. Chretien, I. Arnal, R. H. Wade, Structural changes accompanying GTP hydrolysis in microtubules: information from a slowly hydrolyzable analogue guanylyl-(alpha,beta)-methylene-diphosphonate. J Cell Biol 128, 117–125 (1995).

5. D. Chretien, S. D. Fuller, E. Karsenti, Structure of growing microtubule ends: two-dimensional sheets close into tubes at variable rates. J Cell Biol 129, 1311–1328 (1995).

6. L. M. Rice, E. A. Montabana, D. A. Agard, The lattice as allosteric effector: structural studies of alphabeta- and gamma-tubulin clarify the role of GTP in microtubule assembly. Proc Natl Acad Sci U S A 105, 5378–5383 (2008).

7. D. Bray, Cell movements: from molecules to motility. (Garland Science, 2000).

8. J. Chen, Y. Kanai, N. J. Cowan, N. Hirokawa, Projection domains of MAP2 and tau determine spacings between microtubules in dendrites and axons. Nature 360, 674–677 (1992).

9. T. D. Pollard, W. C. Earnshaw, J. Lippincott-Schwartz, G. Johnson, Cell biology E-book. (Elsevier Health Sciences, 2016).

10. N. Hirokawa, S. Hisanaga, Y. Shiomura, MAP2 is a component of crossbridges between microtubules and neurofilaments in the neuronal cytoskeleton: quick-freeze, deep-etch immunoelectron microscopy and reconstitution studies. J Neurosci 8, 2769–2779 (1988).

11. M. Takeuchi, S. Hisanaga, T. Umeyama, N. Hirokawa, The 72-kDa microtubule-associated protein from porcine brain. J Neurochem 58, 1510–1516 (1992).

12. Conde, A. Caceres, Microtubule assembly, organization and dynamics in axons and dendrites. Nat Rev Neurosci 10, 319–332 (2009).

13. G. Drubin, S. C. Feinstein, E. M. Shooter, M. W. Kirschner, Nerve growth factor-induced neurite outgrowth in PC12 cells involves the coordinate induction of microtubule assembly and assembly-promoting factors. J Cell Biol 101, 1799–1807 (1985).

14. G. Drubin, M. W. Kirschner, Tau protein function in living cells. J Cell Biol 103, 2739–2746 (1986).

15. A. Peters, S. L. Palay, H. d. F. Webster. (Oxford University Press, 1991).

16. S. L. Palay, C. Sotelo, A. Peters, P. M. Orkand, The axon hillock and the initial segment. J Cell Biol 38, 193–201 (1968).

17. M. N. Rasband, The axon initial segment and the maintenance of neuronal polarity. Nat Rev Neurosci 11, 552–562 (2010).

18. X. Li et al., Novel diffusion barrier for axonal retention of Tau in neurons and its failure in neurodegeneration. EMBO J 30, 4825–4837 (2011).

19. N. Drechsel, A. A. Hyman, M. H. Cobb, M. W. Kirschner, Modulation of the dynamic instability of tubulin assembly by the microtubule-associated protein tau. Mol Biol Cell 3, 1141–1154 (1992).

20. N. Gustke, B. Trinczek, J. Biernat, E. M. Mandelkow, E. Mandelkow, Domains of Tau-Protein and Interactions with Microtubules. Biochemistry 33, 9511–9522 (1994).

21. D. Panda, B. L. Goode, S. C. Feinstein, L. Wilson, Kinetic stabilization of microtubule dynamics at steady state by tau and microtubule-binding domains of tau. Biochemistry 34, 11117–11127 (1995).

22. B. Trinczek, J. Biernat, K. Baumann, E. M. Mandelkow, E. Mandelkow, Domains of tau protein, differential phosphorylation, and dynamic instability of microtubules. Mol Biol Cell 6, 1887–1902 (1995).

23. D. Panda, J. C. Samuel, M. Massie, S. C. Feinstein, L. Wilson, Differential regulation of microtubule dynamics by three- and four-repeat tau: implications for the onset of neurodegenerative disease. Proc Natl Acad Sci U S A 100, 9548–9553 (2003).

24. D. W. Cleveland, S.-Y. Hwo, M. W. Kirschner, Purification of tau, a microtubule-associated protein that induces assembly of microtubules from purified tubulin. Journal of Molecular Biology 116, 207–225 (1977).

25. K. S. Kosik, C. L. Joachim, D. J. Selkoe, Microtubule-associated protein tau (tau) is a major antigenic component of paired helical filaments in Alzheimer disease. Proc Natl Acad Sci U S A 83, 4044–4048 (1986).

26. M. Hutton et al., Association of missense and 5’-splice-site mutations in tau with the inherited dementia FTDP-17. Nature 393, 702–705 (1998).

27. A. C. McKee et al., Chronic traumatic encephalopathy in athletes: progressive tauopathy after repetitive head injury. J Neuropathol Exp Neurol 68, 709–735 (2009).

28. Himmler, D. Drechsel, M. W. Kirschner, D. W. Martin, Jr., Tau consists of a set of proteins with repeated C-terminal microtubule-binding domains and variable N-terminal domains. Mol Cell Biol 9, 1381–1388 (1989).

29. K. A. Butner, M. W. Kirschner, Tau protein binds to microtubules through a flexible array of distributed weak sites. Journal of Cell Biology 115, 717–730 (1991).

30. G. Lee, R. L. Neve, K. S. Kosik, The Microtubule Binding Domain of Tau-Protein. Neuron 2, 1615–1624 (1989).

31. B. L. Goode, S. C. Feinstein, Identification of a novel microtubule binding and assembly domain in the developmentally regulated inter-repeat region of tau. J Cell Biol 124, 769–782 (1994).

32. P. Miller, L. Wilson, in Methods in Cell Biology, L. C. J. J. Wilson, Ed. (Elsevier Inc., 2010), vol. 95, pp. 3–15.

33. P. J. Chung et al., Direct force measurements reveal that protein Tau confers short-range attractions and isoform-dependent steric stabilization to microtubules. Proc Natl Acad Sci U S A 112, E6416–6425 (2015).

34. R. Milo, R. Phillips, Cell biology by the numbers. (Garland Science, 2015).

35. P. J. Chung et al., Tau mediates microtubule bundle architectures mimicking fascicles of microtubules found in the axon initial segment. Nat Commun 7, 12278 (2016).

36. Y. Kanai, J. Chen, N. Hirokawa, Microtubule bundling by tau proteins in vivo: analysis of functional domains. Embo J 11, 3953–3961 (1992).

37. E. H. Kellogg et al., Near-atomic model of microtubule-tau interactions. Science 360, 1242–1246 (2018).

38. X. H. Li, J. A. Culver, E. Rhoades, Tau Binds to Multiple Tubulin Dimers with Helical Structure. J Am Chem Soc 137, 9218–9221 (2015).

39. X. H. Li, E. Rhoades, Heterogeneous Tau-Tubulin Complexes Accelerate Microtubule Polymerization. Biophys J 112, 2567–2574 (2017).

40. R. L. Best et al., Tau isoform-specific stabilization of intermediate states during microtubule assembly and disassembly. 294, 12265–12280 (2019).

41. M. A. Ojeda-Lopez et al., Transformation of taxol-stabilized microtubules into inverted tubulin tubules triggered by a tubulin conformation switch. Nat Mater 13, 195–203 (2014).

42. M. T. Gyparaki et al., Tau forms oligomeric complexes on microtubules that are distinct from tau aggregates. Proc Natl Acad Sci U S A 118, (2021).

43. R. Tan et al., Microtubules gate tau condensation to spatially regulate microtubule functions. Nat Cell Biol 21, 1078–1085 (2019).

44. V. Siahaan et al., Kinetically distinct phases of tau on microtubules regulate kinesin motors and severing enzymes. Nat Cell Biol 21, 1086–1092 (2019).

45. J. Needleman et al., Higher-order assembly of microtubules by counterions: from hexagonal bundles to living necklaces. Proc Natl Acad Sci U S A 101, 16099–16103 (2004).

46. G. Beaucage, Approximations Leading to a Unified Exponential/Power-Law Approach to Small-Angle Scattering. Journal of applied crystallography 28, 717–728 (1995).

47. M. C. Choi et al., Human microtubule-associated-protein tau regulates the number of protofilaments in microtubules: a synchrotron x-ray scattering study. Biophys J 97, 519–527 (2009).

48. M. Andreu et al., Low resolution structure of microtubules in solution. Synchrotron X-ray scattering and electron microscopy of taxol-induced microtubules assembled from purified tubulin in comparison with glycerol and MAP-induced microtubules. J Mol Biol 226, 169–184 (1992).

49. H. P. Erickson, W. A. Voter, Polycation-induced assembly of purified tubulin. Proc Natl Acad Sci U S A 73, 2813–2817 (1976).

50. A. Shemesh, A. Ginsburg, Y. Levi-Kalisman, I. Ringel, U. Raviv, Structure, Assembly, and Disassembly of Tubulin Single Rings. Biochemistry 57, 6153–6165 (2018).

51. A. Harada et al., Altered microtubule organization in small-calibre axons of mice lacking tau protein. Nature 369, 488–491 (1994).

52. P. H. S. George-Hyslop, Piecing together Alzheimer’s. Scientific American 283, 76–83 (2000).

53. K. J. Rosenberg, J. L. Ross, H. E. Feinstein, S. C. Feinstein, J. Israelachvili, Complementary dimerization of microtubule-associated tau protein: Implications for microtubule bundling and tau-mediated pathogenesis. Proc Natl Acad Sci U S A 105, 7445–7450 (2008).

54. M. C. Choi et al., Paclitaxel suppresses Tau-mediated microtubule bundling in a concentration-dependent manner. Biochim Biophys Acta Gen Subj 1861, 3456–3463 (2017).

55. G. Fagherazzi, Small angle X-ray scattering edited by O. Glatter and O. Kratky. Acta Crystallographica Section A: Foundations of Crystallography 39, 500–500 (1983).

56. Mylonas et al., Domain conformation of tau protein studied by solution small-angle X-ray scattering. Biochemistry 47, 10345–10353 (2008).

57. E. Kohn et al., Random-coil behavior and the dimensions of chemically unfolded proteins. Proc Natl Acad Sci U S A 101, 12491–12496 (2004).

58. Rubinstein, R. H. Colby, Polymer physics. (Oxford university press New York, 2003), vol. 23.

59. Dehmelt, S. Halpain, The MAP2/Tau family of microtubule-associated proteins. Genome Biol 6, 204 (2005).

60. S. F. B. van Beuningen et al., TRIM46 Controls Neuronal Polarity and Axon Specification by Driving the Formation of Parallel Microtubule Arrays. Neuron 88, 1208–1226 (2015).

61. M. Curcio, F. Bradke, Microtubule Organization in the Axon: TRIM46 Determines the Orientation. Neuron 88, 1072–1074 (2015).

62. M. Harterink et al., TRIM46 Organizes Microtubule Fasciculation in the Axon Initial Segment. J Neurosci 39, 4864–4873 (2019).

63. R. L. Best et al., Expression and isolation of recombinant tau. Methods Cell Biol 141, 3–26 (2017).

64. S. Kar, J. Fan, M. J. Smith, M. Goedert, L. A. Amos, Repeat motifs of tau bind to the insides of microtubules in the absence of taxol. EMBO J 22, 70–77 (2003).

65. V. Makrides, M. R. Massie, S. C. Feinstein, J. Lew, Evidence for two distinct binding sites for tau on microtubules. Proc Natl Acad Sci U S A 101, 6746–6751 (2004).

