## Supplementary material for "Complexes of tubulin oligomers and tau form an intervening network cross-bridging microtubules into bundles": SI Appendix

### <u>Supporting Information (SI) Appendix</u>

**SI Note S1.** *Tubulin and tau purification*

**S1.1** ***Bovine Tubulin Purification***

Crude brain extract was subjected to two cycles of glycerol-free polymerization and depolymerization with centrifugation washes between each step. Resultant MAP-rich microtubules were separated into MAPs and tubulin through a phosphocellulose cationic exchange column. Purified bovine tubulin was purified in PEM50 (50 mM PIPES at pH 6.8, 1 mM EGTA, 1mM $MgSO_4$) with protein concentration between 7 and 14 mg/mL as measured by Braford assay using BSA as a standard.

**S1.2** ***Recombinant Tau Expression and Purification***

Tau was expressed in BL21(DE3) pLacI cells (Invitrogen) with 18 hour incubation in 250 mL of LB media (10 g of tryptone, 5 g of yeast extract, and 10 g of NaCl per liter of DI water) followed by 24 hour incubation in 6 L of auto-induction media (10 g of tryptone, 5 g of yeast extract, 0.5 g of dextrose, 2 g of $\alpha$-$D$-lactose and 5 mL of glycerol per liter of 25mM $NaHPO_4$, 25 mM $KH_2PO_4$, 50 mM $NH_4Cl$, 5 mM $Na_2SO_4$ in DI water). Bacteria were harvested by centrifugation in a Sorvall RC-5B Plus centrifuge at 5000 RPM for 10 minutes maintained between 4 °C and 10 °C. Bacteria resuspended in BRB80 buffer (80 mM PIPES at pH 6.8, 1 mM EGTA and 1 mM $MgSO_4$) were lysed by passing through a French pressure cell press 3 times at >900 PSI, subsequently boiled for 10 min, and then centrifuged at 10,000 RPM for 10 minutes. The supernatant was collected and passed over a phosphocellulose anionic exchange column and eluted with increasing concentration of $(NH_4)_2SO_4$ in BRB80 (up to 1 M). Tau-containing fractions were subsequently pooled and brought to 1.25 M $(NH_4)_2SO_4$, then further purified using hydrophobic interaction column chromatography (HisTrap Phenyl HP, GE Healthcare), eluted with decreasing concentration of $(NH_4)_2SO_4$ in BRB80. Fractions containing pure tau were pooled, then concentrated and buffer-exchanged into PEM50 buffer (50 mM PIPES, 1 mM EGTA, 1 mM $MgSO_4$, brought to pH 6.8 with NaOH) by successive centrifugation cycles using Amicon Ultra-15 Centrifugal Units with MWCO = 10,000 (EMD Millipore, Darmstadt, Germany). Final tau stocks were stored at -80 °C until needed for experiments. Concentration was determined by SDS-PAGE comparison with a tau mass standard, the concentration of which had been established by protein mass spectrometry (1).

**SI Note S2.** *Sample Preparation*

**S2.1 *Whole-mount TEM***

For whole-mount TEM, samples were diluted into warm buffer to 0.2 mg/mL and loaded onto highly stable Formvar carbon-coated copper grids (Ted Pella, Redding, CA), with excess solution wicked with Whatmann paper after 2 minutes. 1% uranyl acetate was added to the grid for 20 seconds and wicked off, and then five drops of Milipore $H_2O$ (18.2 MΩ) were added and wicked off. Grids were allowed to dry for 24 hours at room temperature before imaging.

**S2.2 *Plastic-embedded TEM***

For plastic-embedded TEM, samples were centrifuged to a pellet in microcentrifuge tubes at 9,500 x *g* at 37 °C for 30 min. Supernatant was removed and pellet fixed with 2% glutaraldehyde and 4% tannic acid overnight. The pellet was stained with 0.8% $OsO_4$ in PEM50 buffer for 1 hour and subsequently rinsed four times with PEM50. Another stain of 1% uranyl acetate stain was applied for 1 hour and rinsed with DI water. Fixed and stained pellets were subsequently dehydrated with 25/50/75/100% solutions of acetone in DI water for 15 minutes apiece. Samples were embedded in resin, then embedded in spur plastic and incubated overnight, with resin poured into flat embedding moulds and held at 65 °C for 48 hours and cooled overnight. Plastic-embedded samples were then cut to ~ 50-nm slices with a microtome (Ted Pella, Redding, CA) and transferred to Formvar carbon-coated copper EM grids.

**SI Note S3.** *SAXS Analysis*

Data from 2D scattering images obtained with a Pilatus3 X 1M 2D-detector were azimuthally averaged and subsequently fit to a model containing three terms:

$$I(q) = \iint S(q_r)|F_{MT}(q_r, q_z)|^2 + \iint |F_{Ring}(q_r, q_z)|^2 + BG(q) \qquad (1)$$

where $q_r$ and $q_z$ are wavevectors perpendicular and parallel to the tubular MT axis. Fits were obtained using nonlinear least-squared fitting routines in C-PLOT. The first term in Eq. (1) models the scattering contribution of tubulin in the bundled microtubule state (the structure factor of the MT lattice multiplied by the form factor of a MT and averaged over all orientations in q-space), the second term models the scattering contribution of tubulin in the ring state (form factor of a tubulin ring averaged over all orientations in q-space), and the third term approximates the background scattering contribution including, from (non-ring) unpolymerized tubulin oligomers with small curvature.

The structure factor of the bundled MT state was modelled as the sum of square Lorentzians at every 2D reciprocal lattice vector $q_{hk} = q_{10}(h^2+k^2+hk)^{1/2}$ with amplitude $A_{hk}$ and peak width $\kappa_{hk}$ *(2)*:

$$S(q_r) = \sum_{h,k}[A_{hk}/(\kappa_{hk} + (q_r - q_{10}\sqrt{h^2+k^2+hk})^2)]^2 \qquad (2)$$

The first three Bragg peaks ($q_{10}$, $q_{11}=3^{1/2}q_{10}$, $q_{20}=2q_{10}$) were individually fit, while all other peaks were thereafter fit simultaneously. To limit the number of fitting parameters, all simultaneously fit peaks were assumed to have the same peak width as $\kappa_{20}$ (the highest order individually fit peak). The center-to-center distance between microtubules is given by $a_h = 4\pi/(q_{10}\sqrt{3})$, and the the coherent domain size of the MT lattice (i.e. the hexagonal bundle width) is inversely related to the width of the structure factor peaks and is given by $L_{domain} = 2(\pi \ln 4)^{1/2}/\kappa_{10})$ (3).

The form factor of both the MT and ring states were calculated by modeling them as very long and very short hollow cylinders, respectively, with MT length $L_{MT}$, and ring length $L_{Ring}$ (2, 4):

$$|F_{MT}|^2 \propto \left| [(\sin(q_z L_{MT}/2)/q_r q_z)]\left[(r_{in}+w)J_1\left(q_r(r_{in}+w)\right) - r_{in}J_1(q_r r_{in})\right] \right|^2 \quad (3)$$

$$\left|F_{Ring}\right|^2 = \left| A_{ring}[(\sin(q_z L_{Ring}/2)/q_r q_z)]\left[(r_{in}+w)J_1\left(q_r(r_{in}+w)\right) - r_{in}J_1(q_r r_{in})\right] \right|^2 \quad (4)$$

Here, $J_1$ is the Bessel function of order 1, $r_{in}$ is the ensemble-averaged inner radius of the MT, in Eq. (3), or the ring, in Eq. (4), $w$ is the width of a tubulin monomer (width of MT wall in Eq. (3) or of the ring wall in Eq. (4)) and $A_{ring}$ is the scattering amplitude of the ring state. The average MT length ($L_{MT} \approx 20$ µm) for tau coated MTs (5) is much larger than $1/q_{minimum}$ of the q-range and therefore does not affect the form factor contribution in the measured SAXS range. $L_{Ring}$ was fixed at 49Å to be consistent with electron microscopy data (6).

Any SAXS scattering contribution resulting from neither of these two states (first and second terms in Eq. (1)) was modeled simply as a two layer, unified fit function (7):

$$BG(q) = G\exp\left(\frac{-q^2R_g^2}{3}\right) + B_1\left[\frac{erf\left(\frac{qR_g}{\sqrt{6}}\right)^3}{q}\right]^{-P_1} + \exp\left(\frac{-q^2R_g^2}{3}\right)B_2 q^{-P_2} + BG_0 \quad (5)$$

Where the first term is Guiner's law, the second and third terms are power-law scattering terms which describe the scattering at lengths where $qR_g >> 1$ and $qR_g << 1$, and $BG_0$ is a flat scattering term independent of q. Scattering from the third and fourth term in the equation above dominate the total scattering profile ($I$(q), Eq. 1) at very low-q (<0.005 $A^{-1}$) and high-q (>0.15 $A^{-1}$) respectively, and thus were individually fit within these two domains. Fitting bounds for $R_g$ were determined by fits of the unified scattering function to unpolymerized tau tubulin samples (with scattering contributions from minimal tubulin rings), and was fit simultaneously with the first Bragg as its scattering was most prominent at this length scale ($R_g$=15-20nm). Lastly the fitting parameters of the second term dominate the BG scattering signal for our samples between q=0.03-.09$A^{-1}$, and were thus determined by fitting it to the set of points measured at time $t_0$ where the scattering contribution of the bundled MT state to the raw data was approximately zero (i.e. at the Bessel function minima).

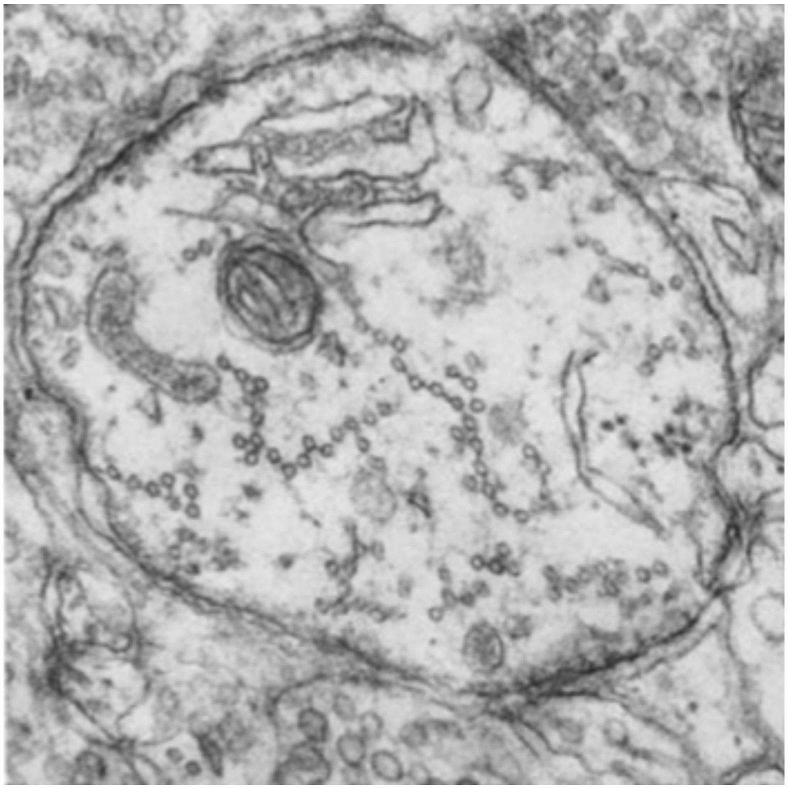

**SI Figure S1. Prior electron microscopy revealed linear microtubule bundles (microtubule fascicles) in vivo in the axon initial segment.** Linear microtubule bundles in the axon initial segment viewed in a transverse section of rat cerebral cortex. X 50,000. Adapted from (8), Fig. 6.

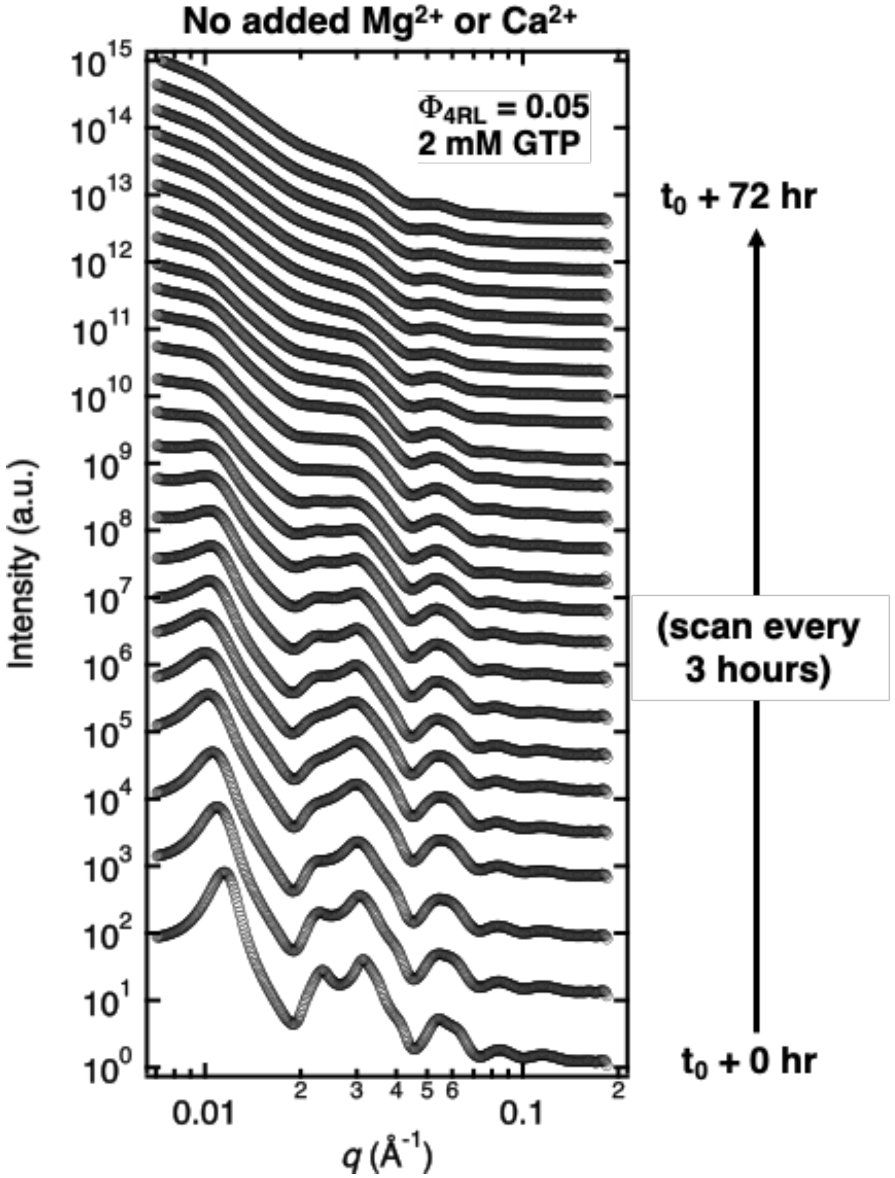


**SI Figure S2. Time- dependent synchrotron SAXS data reveals the stability of the wide-spacing ($B_{ws}$) microtubule bundle state in the absence of added divalent cations.** SAXS profiles are of tubulin/tau/GTP mixtures at 37°C and 4RL-tau to tubulin-dimer molar ratio $\Phi_{4RL}$ = 0.05. $t_0$ corresponds to the short time point after sample preparation when the initial SAXS measurement was performed. Azimuthally averaged synchrotron SAXS data (open circles) with increasing time. SAXS scans are offset for clarity. The $B_{ws}$ was stable over the duration of the experiment (72 hours) and only shows a slow broadening and decrease in intensity of the peaks with increasing time due to ongoing suppressed MT dynamic instability in the presence of tau. Despite gradual MT depolymerization over time, scattering from tubulin rings is absent at all time points.

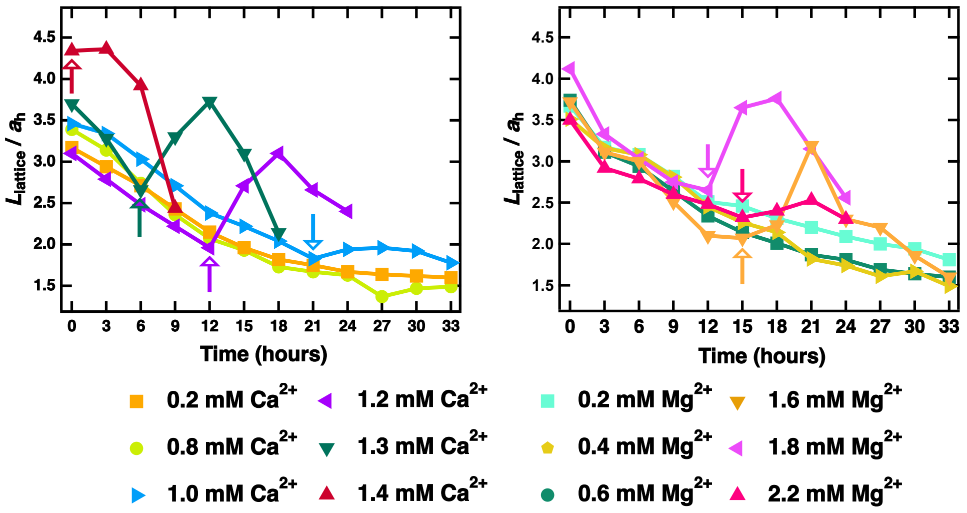


**SI Figure S3. Change in the average domain size of the hexagonal lattice, upon transitioning from the wide-spacing ($B_{ws}$) to the intermediate ($B_{int}$) MT bundle state.**

Time "0" on the x-axis corresponds to the short time point before SAXS data was taken right after sample preparation (referred to as $t_0$ in figures 2,3). Plots of fitted domain size ($L_{domain}$) normalized by the lattice parameter ($a_h$) as a function of time for the SAXS data shown in figures 2 and 3 for the $Ca^{2+}$ (left) and $Mg^{2+}$ (right) series, respectively. Some data points omitted for clarity. Arrows indicate the latest time point the wide-spacing ($B_{ws}$) state is observed as the sample transitions to the intermediate $B_{int}$ state and domain sizes increase.

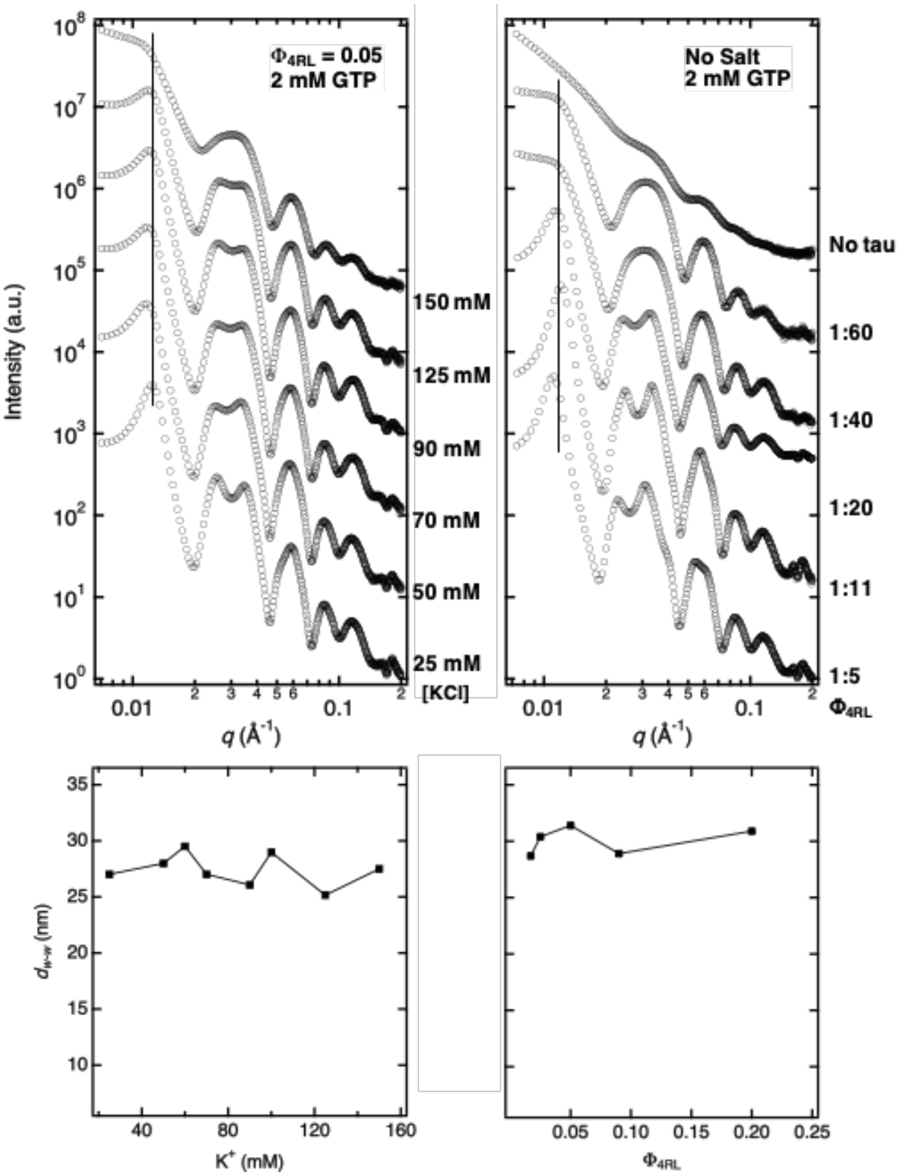


**SI Figure S4. Synchrotron SAXS data reveal the wall-to-wall distance of bundled microtubules is not dependent on KCl concentration or 4RL-Tau to tubulin-dimer molar ratio.** SAXS profiles are of tubulin/tau/GTP mixtures at 37°C and 4RL-tau to tubulin-dimer molar ratio $\Phi_{4RL}$ = 0.05 with increasing KCl concentration (left) and at standard buffer conditions with increasing $\Phi_{4RL}$ (right) at $t_0$. Plots of fitted wall-to-wall distance ($d_{w-w}$) for the corresponding SAXS data highlight the lack of change in $d_{w-w}$ both with increasing KCl concentration and with increasing $\Phi_{4RL}$. (Top) Azimuthally averaged synchrotron SAXS data (open circles) with increasing KCl concentration (left) and decreasing $\Phi_{4RL}$ (right). SAXS scans are offset for clarity. The location of the (1,0) peak, $q_{1,0}$, which is used to measure the center-to-center distance between microtubules is not dependent on KCl concentration or $\Phi_{4RL}$. (Bottom) Plots of fitted wall-to-wall spacings ($d_{w-w}$) as a function of KCl concentration (left) and $\Phi_{4RL}$ (right) of the SAXS data shown above.

**SI Appendix References**

1. D. Panda, J. C. Samuel, M. Massie, S. C. Feinstein, L. Wilson, Differential regulation of microtubule dynamics by three- and four-repeat tau: implications for the onset of neurodegenerative disease. *Proc Natl Acad Sci U S A* **100**, 9548-9553 (2003).
2. D. J. Needleman *et al.*, Higher-order assembly of microtubules by counterions: from hexagonal bundles to living necklaces. *Proc Natl Acad Sci U S A* **101**, 16099-16103 (2004).
3. M. A. Ojeda-Lopez *et al.*, Transformation of taxol-stabilized microtubules into inverted tubulin tubules triggered by a tubulin conformation switch. *Nat Mater* **13**, 195-203 (2014).
4. M. C. Choi *et al.*, Human microtubule-associated-protein tau regulates the number of protofilaments in microtubules: a synchrotron x-ray scattering study. *Biophys J* **97**, 519-527 (2009).
5. R. Brandt, G. Lee, Functional organization of microtubule-associated protein tau. Identification of regions which affect microtubule growth, nucleation, and bundle formation in vitro. *J Biol Chem* **268**, 3414-3419 (1993).
6. J. M. Andreu *et al.*, Low resolution structure of microtubules in solution. Synchrotron X-ray scattering and electron microscopy of taxol-induced microtubules assembled from purified tubulin in comparison with glycerol and MAP-induced microtubules. *J Mol Biol* **226**, 169-184 (1992).
7. G. Beaucage, Approximations Leading to a Unified Exponential/Power-Law Approach to Small-Angle Scattering. *Journal of applied crystallography* **28**, 717-728 (1995).
8. S. L. Palay, C. Sotelo, A. Peters, P. M. Orkand, The axon hillock and the initial segment. *J Cell Biol* **38**, 193-201 (1968).
